# Cargo adaptors use a handhold mechanism to engage with myosin V for organelle transport

**DOI:** 10.1101/2025.03.24.645041

**Authors:** Hye Jee Hahn, Natalya Pashkova, Michael A. Cianfrocco, Lois S. Weisman

## Abstract

Myo2, a myosin V motor, is essential for organelle transport in budding yeast. Its attachment to and detachment from cargo are mediated by adaptor molecules. Vac17, a vacuole-specific adaptor, links Myo2 to Vac8 on the vacuole membrane, and plays a key role in the formation and dissociation of the Myo2-Vac17-Vac8 complex. Using genetics, cryo-electron microscopy and structure prediction, we find that Vac17 interacts with Myo2 through two distinct sites rather than a single interface. Similarly, the peroxisome adapter Inp2 engages two separate regions of Myo2, one of which overlaps with Vac17. These findings support a “handhold” model, in which cargo adaptors occupy multiple sites on the Myo2 tail, enhancing motor-cargo interactions and likely providing additional regulatory control over motor recruitment.

**Summary:** This study provides insights into how cargo adaptors bind myosin V. Genetics, cell-based assays, cryo-EM, and AlphaFold, reveal that the vacuole-specific adaptor uses a handhold mechanism to attach to two areas on the myosin V tail. Moreover, evidence is presented that other adaptors use a similar strategy.

## Introduction

Class V myosin motors form homodimers and mediate the directed transport of intracellular cargoes (Trybus, 2008). These molecular motors are ubiquitously expressed and enable intracellular transport of various types of cargoes along filamentous actin. For example, mammalian myosin Va contributes to the transport of melanosomes in melanocytes (Rogers et al., 1999; Wu et al., 1997), the delivery of insulin-containing secretory granules in pancreatic β-cells (Varadi et al., 2005), and the mobilization of the smooth endoplasmic reticulum in Purkinje neurons (Wagner et al., 2011). Myosin Va also regulates the localization of RNA-protein particles in hippocampal neurons (Nalavadi et al., 2012). Myosin Vb is involved in membrane recycling pathways in polarized epithelial cells (Engevik et al., 2019; Lapierre et al., 2001; Roland et al., 2011), while myosin Vc associates with Weibel-Palade bodies in vascular endothelial cells, where it is critical for their exocytosis (Holthenrich et al., 2022). Thus, myosin V plays a fundamental role in diverse biological processes, including cell-type specific functions and cellular architecture. However, the mechanisms underlying precise motor-cargo interactions are only partially understood.

Myosin V activity is tightly controlled to ensure efficient cargo transport. A key regulatory mechanism involves the release of autoinhibition. In its inactive state, myosin V adopts a folded conformation, where intramolecular interactions between the N-terminal motor domain (head) and C-terminal cargo-binding domain (tail) prevent motor activation. These inhibitory contacts are further reinforced by interfaces formed by the lever arm and coiled-coil domains in the central region of myosin V. Cargo adaptors relieve this inhibition by disrupting these intramolecular linkages, thereby activating myosin V motility (Liu et al., 2006; Niu et al., 2022; Thirumurugan et al., 2006). This highlights their essential role in regulating myosin V function. Mutational analysis in yeast suggest that the autoinhibition mechanism is evolutionarily conserved (Donovan and Bretscher, 2015) and requires cargo adaptors to trigger myosin V activation.

In addition to motor activation, cargo adaptors regulate myosin V by ensuring precise spatial and temporal localization of cargoes. They achieve this by directly linking myosin V to its specific cargo, while serving as a regulatory hub that coordinates cargo attachment and release (Hammer and Sellers, 2011; Weisman, 2006). Vacuole inheritance in budding yeast provides an example of how cargo adaptor regulation controls transport dynamics (Hill et al., 1996; Weisman, 2006). Movement of the vacuole relies on the interactions between yeast myosin V, Myo2, the vacuole-specific adaptor, Vac17, and the vacuole scaffold protein, Vac8. Early in the cell-cycle, a portion of the mother cell vacuole is transferred to the emerging bud. This process is initiated by the availability of Vac17, which is transcribed during G1 and G1/S phases (Spellman et al., 1998), and by phosphorylation of the Vac17 protein (Peng and Weisman, 2008). Vac17 tethers Myo2 to Vac8 (Ishikawa et al., 2003; Kim et al., 2023; Tang et al., 2003; Wang et al., 1998). Once sufficient levels of the Myo2-Vac17-Vac8 complex are formed, the vacuole moves along actin cables toward the bud tip. During this directed movement, additional regulators are recruited to Vac17. Upon the vacuole’s arrival at the bud tip, these regulators terminate transport by ubiquitinating and further phosphorylating Vac17 (Wong et al., 2020; Yau et al., 2014; Yau et al., 2017). These modifications lead to the extraction of Vac17, which is then proteolyzed by the proteasome. The degradation of Vac17 and the subsequent release of Myo2 deposit the vacuole at the bud tip. The released Myo2 can continue to transport other cargoes to the mother-bud neck. Thus, cargo adaptors integrate signals from cellular pathways to regulate Myo2-cargo interactions.

Budding yeast have at least ten additional cargo adaptors for Myo2 (Beach et al., 2000; Beningo et al., 2000; Dunkler et al., 2021; Eves et al., 2012; Fagarasanu et al., 2006; Fortsch et al., 2011; Itoh et al., 2004; Itoh et al., 2002; Jin et al., 2011; Lipatova et al., 2008; Otzen et al., 2012; Tang et al., 2003; Yin et al., 2000). Genetic and structural analyses of the Myo2 tail (Catlett and Weisman, 1998; Ishikawa et al., 2003; Liu et al., 2022; Pashkova et al., 2005; Pashkova et al., 2006; Schott et al., 1999; Tang et al., 2019) have identified critical surface residues involved in interactions with many of these cargo adaptors. Notably, these Myo2 sites often converge at two major regions (Eves et al., 2012), suggesting that cargo recognition is highly coordinated. Investigating how cargo adaptors interact with Myo2 may reveal broader principles that govern actin-based transport systems in higher eukaryotes.

Cargo adaptors for actin-based motors are often structurally flexible, and form transient associations between motors and their cargoes. Since full-length adaptors fail to crystallize with myosin V/Myo2, the available complex structures include only short peptide sequences (Pylypenko et al., 2013; Pylypenko et al., 2016; Tang et al., 2019; Wei et al., 2013). This limitation hinders a comprehensive understanding of the molecular mechanisms underlying myosin V/Myo2-cargo interactions. To overcome this challenge, we combined structural biology with cell-based and computational approaches to map the interaction sites between Vac17 and Myo2.

Here, our study reveals that Vac17 interacts with Myo2 through two distinct sites rather than a single site. We identify a Vac17 region that forms a second interface. This N-terminal region of Vac17, now termed Vac17(H), is predicted to adopt a three-helix bundle (h1-3). The presence of both Vac17(H) and its canonical Myo2-binding domain, Vac17(MBD), is necessary for proper Myo2-mediated vacuole movement. Moreover, the structure shows that Vac17(H) and Vac17(MBD) bind to opposite sides of the Myo2 tail. This suggests that Vac17 wraps around the Myo2 tail and binds in a “handhold” mechanism. Additionally, we show that the peroxisome adaptor Inp2 also uses two sites on Myo2. These findings predict that multiple cargo adaptors employ handhold mechanisms to engage Myo2. The binding of adaptors at two sites likely stabilizes the motor-cargo association, and provides additional sites for regulation for the attachment and release of cargoes.

## Results

### Vac17(H) contributes to vacuole inheritance

To probe how Vac17 functions with Myo2, we utilized the *myo2-N1304S* mutant, which is defective in its interaction with Vac17 (Catlett et al., 2000; Ishikawa et al., 2003; Pashkova et al., 2006), and screened for compensatory mutations in Vac17 that could restore vacuole inheritance. We performed random PCR mutagenesis across the entire Vac17 open reading frame to ensure that all regions were included. Eleven suppressor mutations were identified, however, only three mutations (T110P, R126S and I140V) reside within the canonical Myo2-binding domain, Vac17(MBD) (**Fig. 1 A; Table S1**). Notably, many of the suppressor mutations were located upstream of the Vac17(MBD), suggesting that this region, referred to as Vac17(H), also plays a role in its interactions with Myo2.

**Figure 1.**
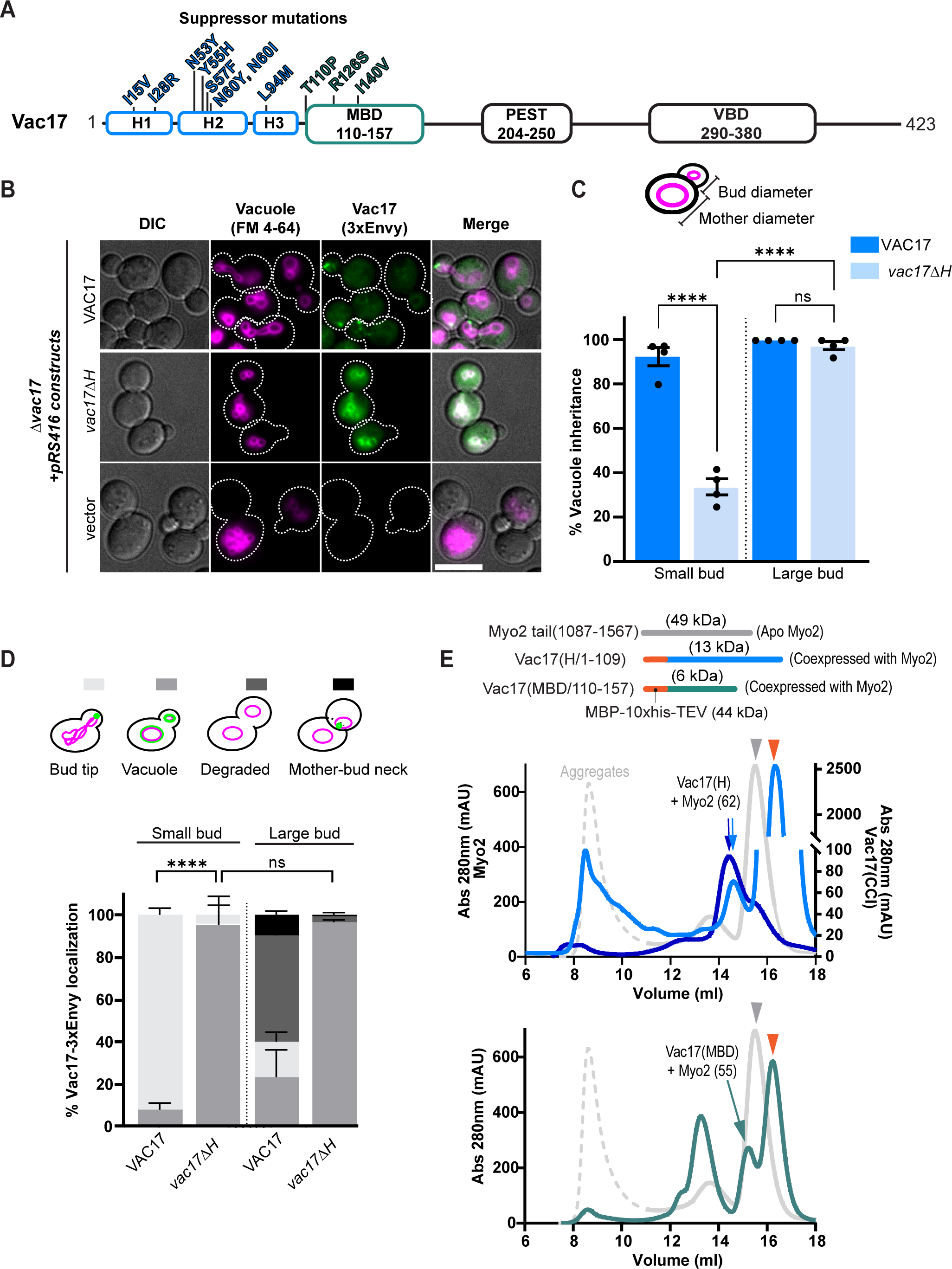
The N-terminal region of Vac17 is important for vacuole inheritance and binds to Myo2. **(A)** Schematic of Vac17. Indicated residues are suppressors identified in a mutagenesis screen that restore vacuole inheritance in the *myo2(N1304S)* mutant. h1-3 = helix 1-3 predicted by all AlphaFold models to adopt an antiparallel 3-helix bundle (H); MBD = Myo2-binding domain; VBD = Vac8-binding domain. **(B)** Vacuole inheritance in *Δvac17* cells transformed with a low-copy plasmid that expresses the indicated Vac17-3xEnvy constructs (green) or empty vector. Vacuole membranes (magenta) were pulse-chase labeled with FM 4-64 and incubated for one doubling time prior to imaging. Dashed lines indicate the cell perimeters. Scale bar = 5 μm. **(C)** Quantification of vacuole inheritance. Scoring was based on the presence or absence of labelled vacuoles at the indicated bud sizes (n=4 experiments; 30 ≥ cells per n for each bud size). Bud size was determined based on the ratio of the diameters of the bud cell to the mother cell. Values ≤ 0.3 was classified as small. Significance was calculated using one-way ANOVA and Tukey’s multiple comparisons; **** P < 0.0001; ns = not significant. Error bars indicate SD. **(D)** Vac17 localization relative to the vacuole in cells where vacuole inheritance occurred (n=4; 30 ≥ cells per n for each bud size). Statistical significance determined by two-way ANOVA and Tukey’s multiple-comparisons test. **** P < 0.0001; ns = not significant. Error bars are SEM. **(E)** Size-exclusion chromatography traces of apo Myo2 tail (gray) alone or co-expressed with N-terminally MBP-fused Vac17 peptides: Vac17(H) (lighter blue; top panel) and Vac17(MBD) (green; bottom panel). The orange arrowhead indicates free maltose-binding protein (MBP) (44 kD), which was cleaved from the Vac17 peptides using TEV protease, following MBP affinity purification. In a separate experiment, Myo2 tail co-expressed with MBP-fused Vac17(H) was first pulled down using amylose resin, treated with TEV protease, and subsequently pulled down with Strep-Tactin to isolate Myo2. Subsequent gel filtration confirms that Vac17 associates with Myo2 (dark blue).

To investigate the role of Vac17(H), we first examined whether this N-terminal sequence forms a coiled-coil, as previously suggested (Tang et al., 2003). Notably, 2D and 3D structural analyses revealed that Vac17(1-109) adopts a three-helix bundle. Thus we have renamed this region Vac17(H) (**Fig. 1 A**). Next, we deleted residues 18-108 and assessed the impact of the *vac17ΔH* mutant on vacuole inheritance. When vacuole inheritance is impaired, new buds form vacuoles through *de novo* synthesis (Jin and Weisman, 2015). To differentiate between inherited and newly synthesized vacuoles, we used pulse-chase labeling of the vacuoles with FM 4-64 (Vida and Emr, 1995) (**Fig. 1 B**). In wild-type cells, vacuole inheritance typically occurs during bud emergence or in small buds. However, in the *vac17ΔH* mutant, approximately 67% of cells with small buds lacked vacuoles, whereas, in cells with large buds, similar to wild-type, approximately 97% of buds contained vacuoles (**Fig. 1 C**). These results indicate that the *vac17ΔH* mutant exhibits a delay in vacuole inheritance.

The localization of Vac17 relative to the vacuole and sites of polarized growth—at the bud tip or the mother-bud neck where Myo2 is located—is another way to measure Vac17 function. In wild-type cells, at the onset of vacuole inheritance, Vac17 accumulates as puncta on a portion of the vacuole near sites of polarized growth. Once the vacuole reaches the bud tip, Vac17 is targeted for degradation, leading to the loss of its signal in cells with large buds (Peng and Weisman, 2008; Wong et al., 2020; Yau et al., 2014; Yau et al., 2017). In contrast, in the *vac17ΔH* mutant, Vac17 failed to form puncta and instead was observed throughout the vacuole membrane in both mother and bud cells (**Fig. 1 D**). This altered localization resembles that seen in Myo2 point mutants that are specifically defective in Vac17 interaction (Eves et al., 2012; Peng and Weisman, 2008). That vacuoles are eventually inherited in large buds suggests that the *vac17ΔH* mutant retains partial association with Myo2. Importantly, the *vac17ΔH* mutant did not alter Myo2 localization (**Fig. S1 A**). Additionally, the Vac8 signal around the vacuole remained unaffected in the *vac17ΔH* mutant (**Fig. S1 B**). The colocalization of the *vac17ΔH* mutant with Vac8 indicates that Vac17(H) is not required for the interaction between Vac17 and Vac8 (**Fig. S1 C**). However, in the absence of Vac8, the *vac17ΔH* mutant becomes cytosolic. While Vac17(MBD) remains essential for vacuole inheritance, (**Fig. S1 D** and (Ishikawa et al., 2003)), Vac17(H), through its interactions with Myo2, facilitates the movement of vacuoles into small buds.

### Vac17(H) and Vac17(MBD) each directly binds to the Myo2 tail *in vitro*

Based on the Vac17 mutant data, we hypothesized that Vac17(H) directly binds Myo2. To test this, we co-expressed and purified recombinant peptides of Vac17(H) or Vac17(MBD) with the Myo2 tail. Consistent with previous biochemical studies, Vac17(MBD) bound to the Myo2 tail (**Fig. 1 E, Fig. S1 E**; (Eves et al., 2012; Liu et al., 2022; Tang et al., 2019). Notably, we observed that the Vac17(H) peptide exhibited affinity for the Myo2 tail. To confirm complex formation, we performed a reciprocal pull-down of Vac17 followed by Myo2. Subsequent gel filtration analysis (darker blue) revealed a peak corresponding to the expected Vac17(H)-Myo2 complex (lighter blue), verifying their interaction. These findings suggest that the partial inheritance defect observed in the *vac17ΔH* mutant results from a reduced ability of Vac17 to bind Myo2.

### Vac17(H) affects the timing of initial vacuole movement into small buds

The above biochemical studies suggested that Vac17(H) directly binds to Myo2, and likely enhances the ability of Myo2 to transport the vacuole into the bud. To assess how Vac17(H) impacts vacuole movement, we performed live-cell imaging in a strain with the vacuole marker, mCherry-Vph1 expressed from its endogenous locus, and transformed with plasmids that express either wild-type or the *vac17ΔH* mutant (**Fig. 2, A-D**). Cells were arrested in G1 phase using alpha-factor and released into fresh media. During this period, cells were transferred into a microfluidics plate, where they were continuously perfused with fresh media. We noted that the Vac17 levels were elevated near the leading edge of the vacuole in synchronized cells. Thus, to avoid the potential effects from excess Vac17 accumulation on the rate of vacuole movement, cells were allowed to complete one doubling cycle prior to time-lapse imaging during the second cycle.

**Figure 2.**
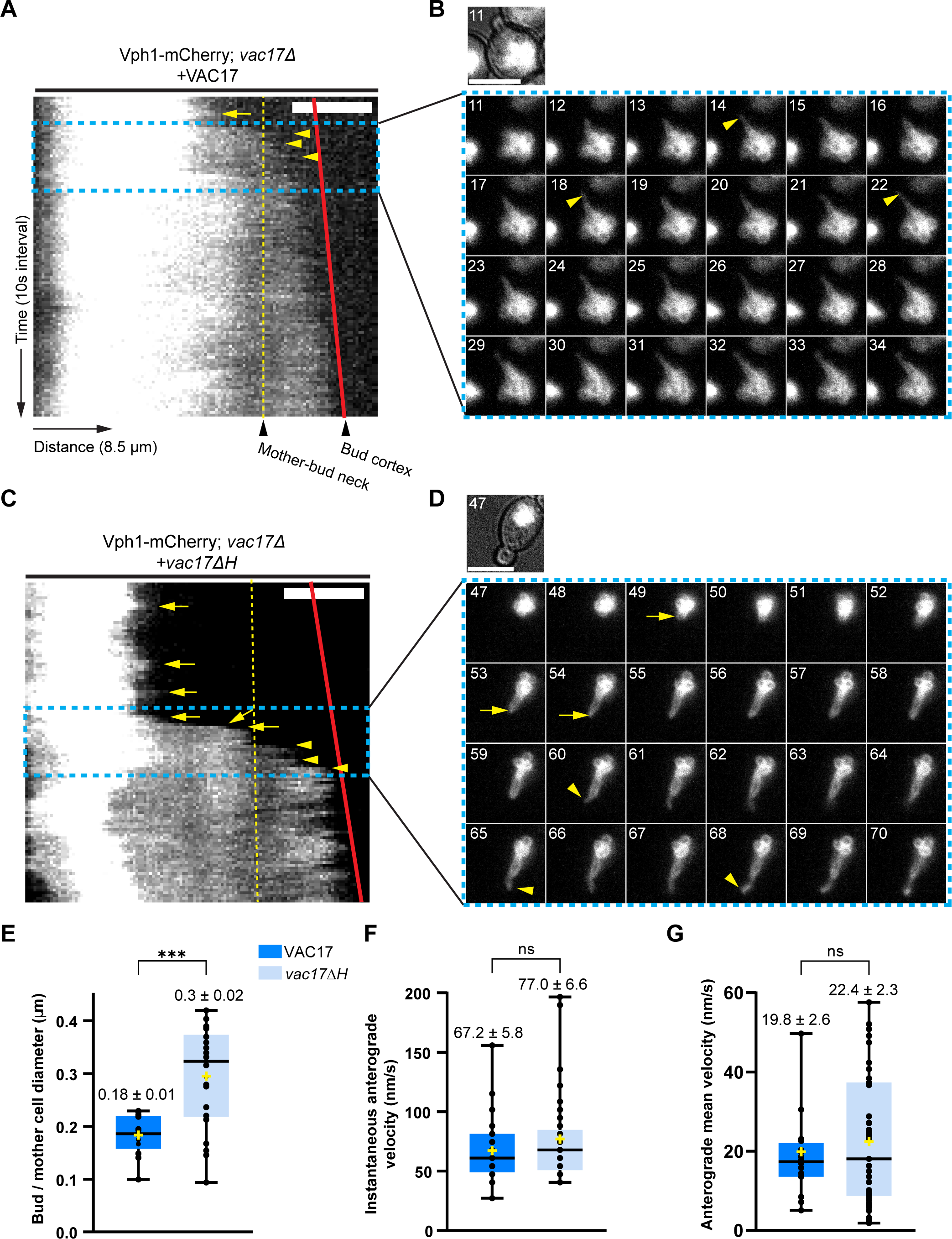
*vac17ΔH* cells show a delay in the initiation of vacuole inheritance. **(A, C)** Kymographs of vacuole dynamics in wild-type and *vac17ΔH* cells. Images were taken on a single z-plane to reduce photobleaching. Y-axis represents time (10 seconds per frame), and X-axis is the directionality of vacuole movement along the segmented path of 8.5μm (0.06788 μm/pixel). Vacuoles are labelled with endogenously expressed Vph1-mCherry, which localizes to the vacuole. Dashed lines (yellow) indicate the mother-bud neck; solid lines (red) represent the growing bud tip. Yellow arrows indicate vacuole anterograde events prior to crossing the mother-bud neck. Yellow arrowheads indicate anterograde movements following vacuole inheritance. Only cells in which the bud had not yet received a vacuole at the start of imaging were included in the following quantifications. **(B, D)** Montage of a time-series of cells undergoing vacuole inheritance. The montage corresponds to the blue box regions of the kymographs. **(E)** Ratio of the bud and mother cell size when the vacuole first crossed the mother bud neck (n=13 and n=22 for wild-type and *vac17ΔH*, respectively). **(F)** Instantaneous velocities of the vacuole movement (n=24 and n=33 for wild-type and *vac17ΔH*, respectively). **(G)** Rate of all anterograde vacuole movements, up until the vacuole reached the bud cortex (n=20 and n=51 for wild-type and *vac17ΔH*, respectively). Significance was determined using unpaired, two-tailed t-test. Data represented as box plots. Colored areas indicate the population encompassing the 1st to 3^rd^ quartile, with the median indicated. The whiskers represent the spread of the data. *** P < 0.001; ns = not significant. Challenges that impacted quantification: 1) multiple movements of the vacuole that were not directional, 2) the distances between the edge of the mother vacuole to the small bud tip was as short as 3.4 μm for wild-type and 5.9 μm for the mutant, and 3) the growing small buds did not always remain in the same plane of focus as the mother cell.

To quantify vacuole movement across the mother-bud neck, we performed kymograph analysis of the mCherry-Vph1 vacuole signal from cells with a bud that was initially devoid of the vacuole (**Fig. 2, A and C**). The red lines on the kymographs represent the growing bud tip, and the yellow dashed lines indicate the mother-bud neck. We also created a montage of time-lapse images from representative cells, including the moment when the vacuole first crosses the mother-bud neck. These image sequences are marked by the blue dashed box in the kymographs (**Fig. 2, B and D**). Consistent with earlier observations, in wild-type cells, when the vacuole crosses the mother-bud neck, the ratio of the bud diameter relative to the mother cell is 0.18. However, in *vac17ΔH* cells, vacuole inheritance is delayed (**Fig. S2, A-C**) resulting in a bud diameter:mother diameter ratio of 0.3 (**Fig. 2 E**).

To determine whether the Myo2 motility is impaired, we calculated the instantaneous velocity of the vacuole in each cell, irrespective of the starting and ending position (**Fig. 2 F**), and found that there was no significant difference (67 µm/s vs. 77 µm/s for wild-type and *vac17ΔH*, respectively). In a second analysis, we analyzed the rates of all anterograde movements per cell, until the first time the signal reached the bud cortex (**Fig. 2 G**). Again, these anterograde velocities in wild-type (19.8 µm/s) vs. the *vac17ΔH* mutant (22.4 µm/s) cells were not significantly different. Given the 10-second interval between image acquisitions, some velocities may be faster than reported. We note that these measured rates are rough approximations. Thus, while Myo2 velocity appears unaffected, the timing of vacuole transport is delayed in the *vac17ΔH* mutant.

Although vacuole inheritance eventually occurs in *vac17ΔH* cells, we observed that unlike wild-type cells, the Vac17 signal persistently localizes around the entire vacuole. Moreover, western blot analysis showed that Vac17 expression was elevated four-fold compared to wild-type (**Fig. S2 D**). Since an intact Myo2-Vac17-Vac8 complex is required for Vac17 degradation (Tang et al., 2003; Yau et al., 2014; Yau et al., 2017), the increased steady-state levels of Vac17 in *vac17ΔH* cells further suggests that this mutant is defective in its interaction with Myo2. We postulate that the excess availability of Vac17 in the *vac17ΔH* mutant cells eventually recruit enough Myo2 motors to initiate vacuole transport.

### Vac17(MBD) binds to the Myo2 tail as an extended, mostly unstructured peptide

Earlier studies were unable to crystallize Myo2 bound to high-affinity peptides derived from the Vac17(MBD) sequence (Liu et al., 2022), leading us to consider cryo-EM. Given the importance of Vac17(H) alongside Vac17(MBD), we set out to characterize how the Vac17(H+MBD) region associates with Myo2 at the molecular level. We purified a recombinant complex comprising co-expressed Myo2 tail and the Vac17(H+MBD) peptide. Gel filtration analysis of the complex showed that Vac17(H+MBD) and Myo2 tail co-eluted as a single peak (**Fig. S3 A**). Vitrification of the sample and subsequent analysis using 2D class averages revealed high-quality, monodisperse particles, indicating that the complex was amenable for structure determination by cryo-EM (**Fig. S3, B-E**).

After collecting and analyzing a dataset for the Vac17(H+MBD)-Myo2 tail, we determined a cryo-EM structure at an overall resolution of ∼6 Å (**Fig. 3 A**). Docking and comparison of this reconstruction with the previously solved crystal structure of the apo Myo2 tail (PDB: 2F6H; (Pashkova et al., 2006) revealed a similar overall architecture (**Fig. S3 F**). Notably, the reconstruction revealed an additional, unoccupied density. Given its proximity to Myo2 residues D1297 and N1304, which are critical for vacuole inheritance, we assigned this density to Vac17 (**Fig. S4 A**; Eves et al., 2012).

**Figure 3.**
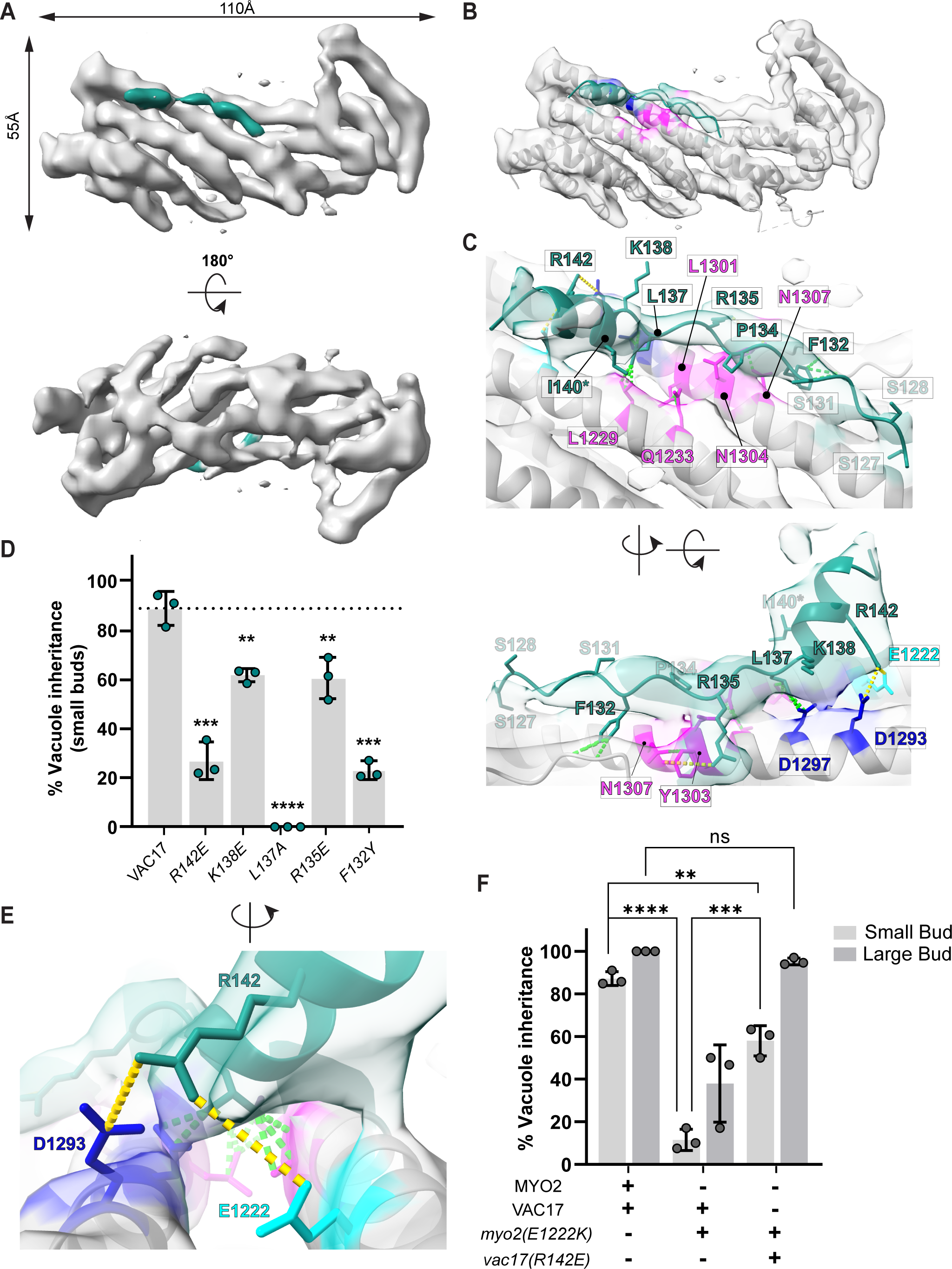
Vac17(MBD) contains an unstructured peptide that extends along the Myo2 tail. **(A)** Cryo-EM reconstruction of the Vac17 peptide (green) bound to the Myo2 tail (gray) **(B)** AlphaFold model of the Vac17(1-157) and Myo2 tail complex was trimmed to fit into the density and refined in real-space using Phenix. Vac17(MBD) density (green) is located near the Myo2 helix (H6), which was previously characterized to contain residues critical for vacuole inheritance (dark blue) and for the inheritance of both vacuole and mitochondria (pink) (Eves et al., 2012; Pashkova et al., 2006). **(C)** Vac17 residues predicted to form contacts with known Myo2 residues. Close contacts are shown in green dashed lines, and interactions over longer distances of 3.8-5 Å are in yellow lines. An asterisk represents one of the Vac17 suppressors, I140V, identified in the screen. This further supports the hypothesis that this region of Vac17 directly interacts with Myo2. **(D)** Vacuole inheritance in small buds of indicated Vac17 mutants. Also, note that in the *vac17(L137A)* mutant, only 6% of the large buds had a vacuole, but in the other mutants reported here, vacuoles were present in large buds (not shown). The significance was determined by unpaired student t-test. Error bars are SD. **** P < 0.0001; *** P < 0.001; ** P < 0.005; ns = not significant. **(E)** Close up view showing predicted interaction of Vac17(R142) with Myo2(E1293) and Myo2(E1222). **(F)** Charge reversal mutations of *vac17(R142E)* and *myo2(E1222K)* partially restore vacuole inheritance. (n=3; ≥ 35 cells per n for each bud size). Significance was determined using one-way ANOVA with multiple-comparisons test. **** P < 0.0001; *** P <0.001; ** P < 0.01; * P < 0.1; ns = not significant.

To characterize the Vac17-Myo2 interaction at the amino acid level, we used the COSMIC^2^ server (Cianfrocco, 2017) to perform AlphaFold2-Multimer structure predictions (Evans et al., 2022; Jumper et al., 2021; Mirdita et al., 2022) of the Vac17(H+MBD) and Myo2 tail complex (**Fig. 3 B**). The predicted models closely recapitulated the atomic structure of the apo Myo2 tail (**Fig. S3 F**). Importantly, the structure of the Vac17(MBD) sequence (residues 127-147) aligned well the putative cryo-EM density for Vac17 (**Fig. 3 C**; Eves et al., 2012). Within this region, Vac17 residues 127-137 formed a disordered loop extending along the Myo2 helix, while residues 138-147 adopted a short helix directed away from Myo2 with minimal apparent contact. Even in the absence of high-resolution cryo-EM data, integrating predictive modeling with experimental density can enable the identification of distinct structural features, offering valuable insights into the flexibility and organization within the complex.

To determine the orientation of Vac17 on the Myo2 tail, we focused on Vac17(R142), a polar residue near the interface that is predicted to interact with Myo2(E1222) (**Fig. 3, D-F**). We hypothesized that these residues form electrostatic interactions, and thus performed charge reversal mutagenesis to assess whether restoring charge complementarity could rescue function. The *vac17(R142E)* mutant impaired vacuole inheritance in small buds (**Fig. 3 D**), consistent with predictions-based isothermal titration calorimetry (ITC) study, which showed a complete loss of affinity between *vac17(R142E)* and the Myo2 tail (Liu et al., 2022). Similarly, the *myo2(E1222K)* mutant displayed defect in inheritance (**Fig. 3, E and F**). Importantly, co-expression of *Vac17(R142E)* and *myo2(E1222K),* partially rescued vacuole inheritance, supporting the proposed orientation of Vac17 binding to Myo2. However, since the *vac17(R142E)* mutant retains partial function *in vivo* and large buds still inherit the vacuole, this suggests that additional interactions are involved.

To further assess how Vac17(MBD) binds to Myo2, we introduced additional mutations within the region of Vac17 identified from the cryo-EM map, and found that hydrophobic interactions play a critical role (**Fig. 3 D**). Most notably, a conservative substitution *vac17(L137A)* abolished vacuole inheritance, consistent with the ITC study showing that *vac17(L137Q)* disrupts Myo2 binding (Liu et al., 2022). Vac17(L137) interacts with Myo2(L1229 and L1301), which form a hydrophobic groove (**Fig. S4, B and C**). This groove is also critical for recognition by Mmr1(L410) for mitochondrial transport (Tang et al., 2019). In contrast, Vac17 residues (R142, K138, and R135), which could mediate multiple polar interactions with Myo2 surrounding the hydrophobic patch, resulted in only partial inheritance defects (**Fig. 3, C and D**). Interestingly, Vac17(I140), a suppressor residue, lies within a helix that does not directly contact Myo2, but may induce local structural adjustments that restore contact with the *myo2(N1304S)* mutant. Collectively, *in vivo* analyses of these mutants combined with the cryo-EM structure, reveals that the Vac17(MBD) and Myo2 tail interaction is primarily driven by hydrophobic contacts, with additional stability provided by polar interactions.

### Conserved features provide insights into yeast and mammalian myosin V interactions with cargo adaptors

To investigate the general principles governing interactions of myosin V and cargo adaptors that bind in the Vac17(MBD) region, we compared our Vac17-Myo2 structure with the Mmr1-Myo2 complex (PDB: 6IXP; Tang et al., 2019; **Fig. 4, A and B**). Previous genetic studies demonstrated that Vac17 and Mmr1 bind to overlapping and distinct sites on Myo2 (Eves et al., 2012). Docking the Mmr1-Myo2 structure revealed that Mmr1 partially occupies the Vac17 density, confirming their shared binding region (**Fig S4. D**). Although Vac17 and Mmr1 adopt similar structural motifs, they bind in opposite orientations. A key hydrophobic residue, Vac17(L137), aligns with Mmr1(L410) to recognize an overlapping site on Myo2 (**Fig. 4 F**). In addition, the positively charged Vac17(R142) interacts with Myo2(E1222), a residue specific to the Vac17-binding region, whereas Mmr1(R409) forms salt bridges with Myo2(E1293) (Tang et al., 2019). Notably, Myo2(E1293), which is situated directly across from Myo2(E1222) in the neighboring helix, was previously thought to be exclusive to Vac17 binding.

**Figure 4.**
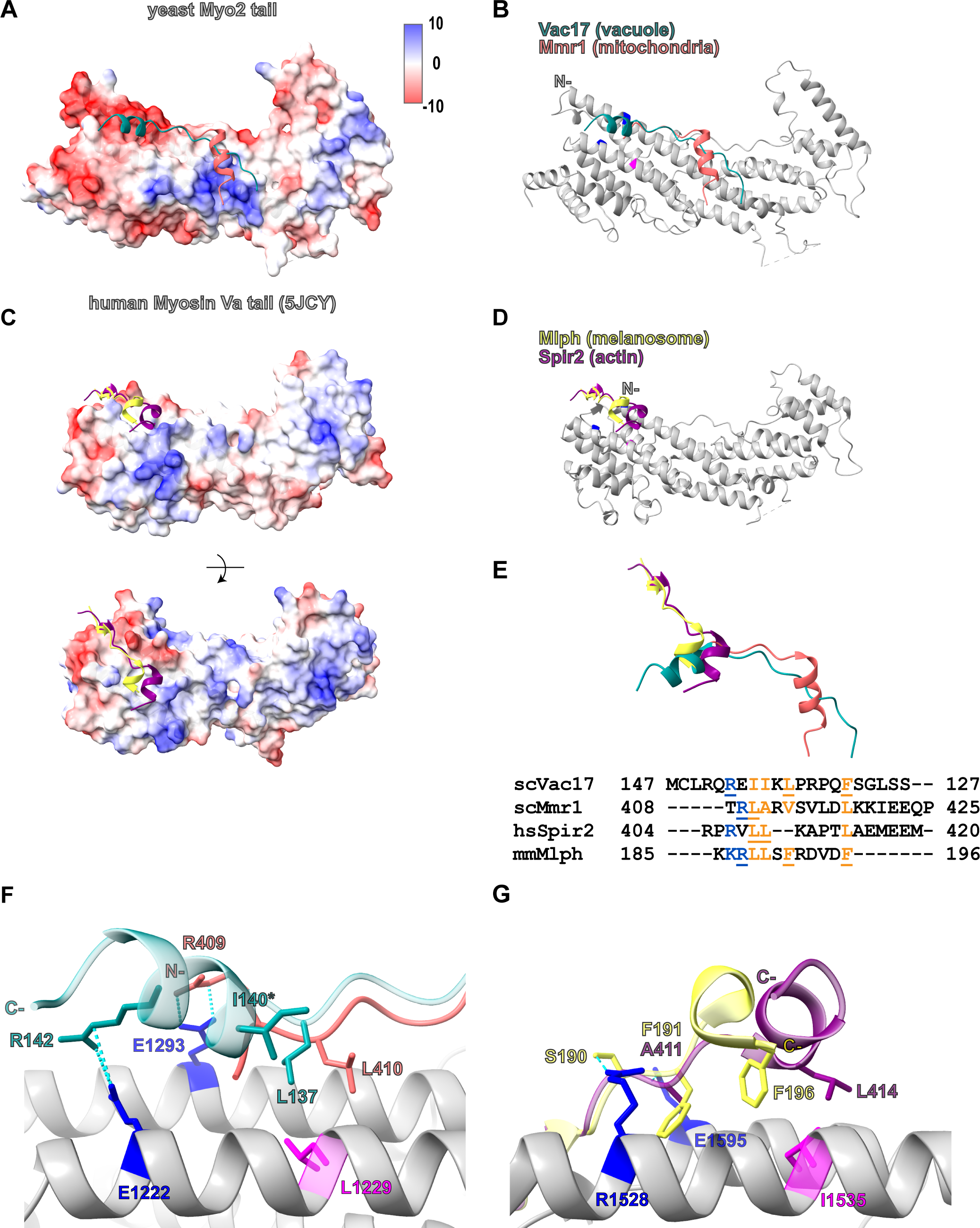
Structural analysis of mammalian myosin Va and yeast Myo2 with their cargo adaptors reveals a common mechanism of interaction. **(A, C)** Electrostatic surface potentials for the yeast and mammalian myosin Va tails, respectively. **(B, D)** Adaptor peptides each interacts with the longest helix of the myosin V tail. **(E)** Ribbon representation illustrates that adaptor peptides adopt similar folds and structurally overlap when aligned to the longest myosin V helix (not shown). Sequence alignment reveals common features within the adaptors that are important for myosin V recognition. Orange indicates hydrophobic residues, and blue represents positively charged residues. Underlined residues are critical for interaction (Pylypenko et al., 2013; Pylypenko et al., 2016; Wei et al., 2013; Tang et al., 2019; this study). Note that Vac17 is the only adaptor that interacts with myosin V in an anti-parallel orientation. **(F)** Vac17(L137) and Mmr1(410), each forms hydrophobic interactions with Myo2(L1229). Hydrophobic contacts are further stabilized by ionic interactions between Vac17(142)-Myo2(E1222) and Mmr1(R409)-Myo2(E1293). Asterisk indicates the suppressor of Vac17(I140), which is spatially proximal to Myo2(L1229). Cyan dashes indicate bond distances between 2.5-5 Å. **(G)** Mlph(F196) interacts with myosin Va(I1535), a key residue mediating hydrophobic contacts (Wei et al., 2013; Pylypenko et al., 2013). Similarly, Spir2(L414) is predicted to form hydrophobic interactions with myosin Va(I1535). In addition, hydrogen bonds are formed between myosin Va(R1528) and Mlph(S190), between myosin Va(E1595) and Mlph(F191), and between myosin Va(E1595)-Spir2(A411). These polar interactions are analogous to the charged contacts between Myo2(E1222) and Vac17(R142) and between Myo2(E1293) and Mmr1(R409). Blue dashed lines represent bond lengths of 2.5-5 Å, and orange dashed lines indicate predicted interactions.

Our analysis further reveals that the Vac17 and Mmr1 binding sites on Myo2 overlap more extensively than previously recognized. *In vitro*, an Mmr1(398-430) peptide with an R409E mutation abolishes binding to Myo2(E1293) (Tang et al., 2019). However, *in vivo* the *Mmr1(R409E)* mutation only partially disrupts mitochondrial inheritance (Nayef et al., 2024), and the *myo2(E1293K)* mutation has no effect on mitochondrial inheritance (Eves et al., 2012). These results suggest that while Mmr1(R409) directly contacts Myo2(E1293), additional charged residues nearby can also interact with Mmr1. Importantly, these adaptors engage Myo2 through multiple interaction sites. Notably, the regions specific to Mmr1 and Vac17 recognition harbor opposite charge potentials around their overlapping hydrophobic patch (**Fig. 4 A**), which likely contributes to the precise positioning of each adaptor.

Building on these new insights, we examined published structures of the mammalian myosin Va tail bound to its cargo adaptors, melanophilin (Mlph) and Spire2 (Spir2) (Pylypenko et al., 2013; Pylypenko et al., 2016; Wei et al., 2013) (**Fig. 4, C and D**). Strikingly, despite significant sequence divergence, the Mlph, Spir2, Vac17 and Mmr1 adaptors converge on a key myosin V/Myo2 cargo recognition site and form interactions that are conserved (**Fig. 4, E**). A key residue Mlph(F196) (Pylypenko et al., 2013; Pylypenko et al., 2016), forms hydrophobic contacts with myosin Va(I1535), which is analogous to the interactions of Vac17(L137) and Mmr1(L409) with Myo2(L1229) (**Fig. 4, F and G**). Interestingly, Spir2(L414) occupies a similar position to Vac17(L137) and Mmr1(L410) and may interact with myosin V(I1535), although this interaction has not yet been tested. Additionally, residues in myosin Va(R1528 and E1595)—analogous to Myo2(E1222 and E1293)—form hydrogen bonds with Mlph(S190 and F191) and Spir2(A411), consistent with the electrostatic interactions observed between Myo2(E1222 and E1293) and Vac17(R142) and Mmr1(R409).

Note that Vac17 uniquely binds in an inverted orientation relative to the longest helix of the motor tail. Thus, 3D structural information was required to discover these conserved binding motifs. Together, these findings reveal that a central hydrophobic core is supported by adjacent polar interactions, underscoring conserved principles in myosin V adaptor recognition.

### Vac17 binds two distinct sites on the Myo2 tail

Due to the absence of an obvious density for Vac17(H), the above reconstruction focused Vac17(MBD) and Myo2. Notably, 2D class averages showed a “fuzzy” density located on the opposite end from Vac17(MBD) (**Fig. 5 A**). To resolve this density, we used a combination of signal subtraction and focused classification on the region showing the fuzzy density from the class averages (**Fig. S5**). Alignment-free 3D classification of these subtracted particles revealed several classes with additional density. By selecting the class with the most continuous density and reverting to un-subtracted particles, we obtained a 3D reconstruction revealing the presence of additional density on the Myo2 tail (**Fig. S5**). To rule out masking artifacts, we performed another round of 2D classification, *ab initio* reconstruction, and 3D refinement, which confirmed the second density on the Myo2 tail.

**Figure 5.**
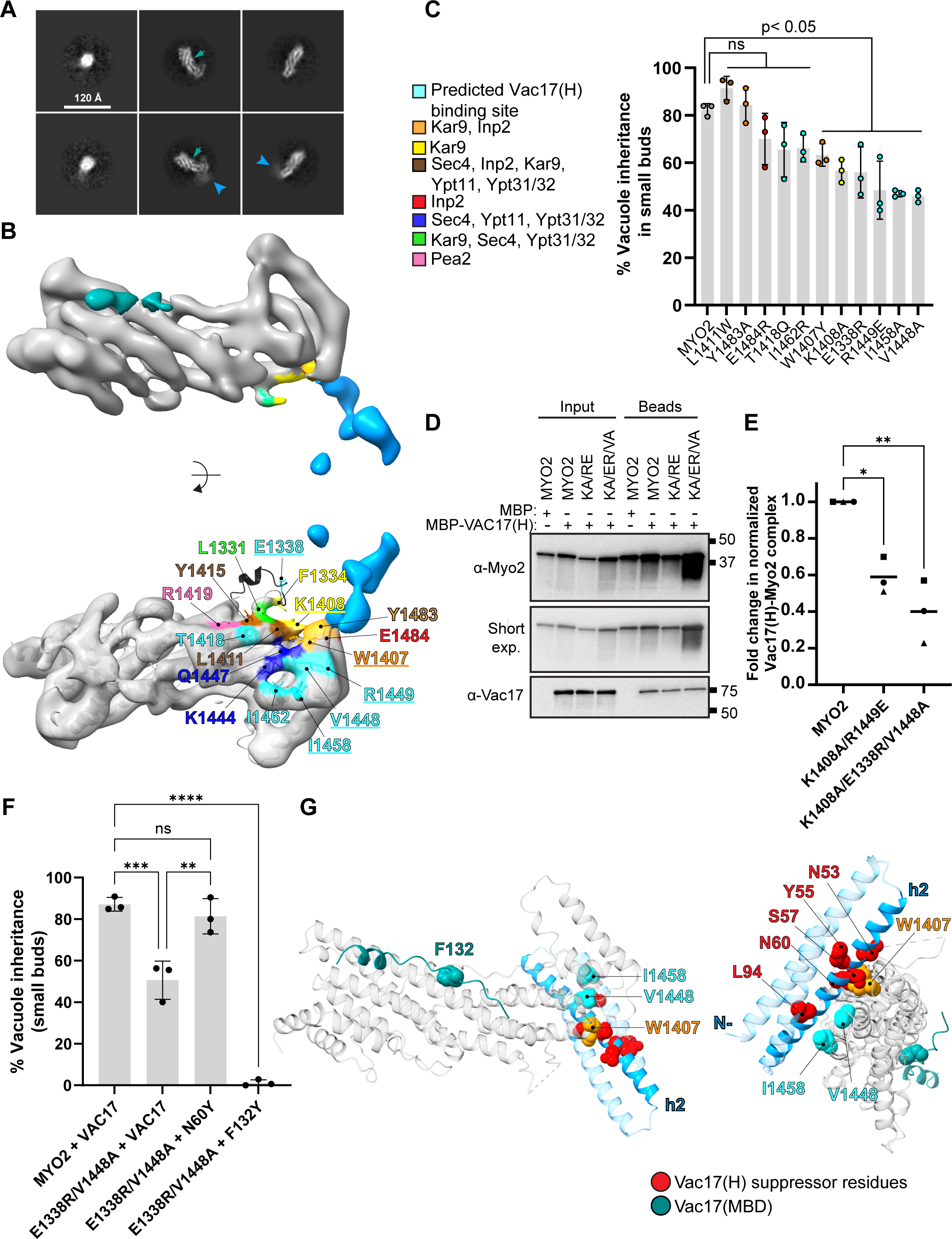
Identification of a second Vac17-Myo2 interface. **(A)** 2D class averages of Vac17-Myo2 without or with the fuzzy density (blue arrowhead). These were observed in the same dataset. Location of the Vac17(MBD) is shown in green arrowhead. **(B)** A second reconstruction of the Vac17-Myo2 interaction includes the additional density (blue) obtained through particle subtraction. Focused alignment-free 3D classification was performed on the subtracted particles. The additional Vac17 density is located near the Myo2 region crucial for transporting various cargoes, including peroxisomes, secretory vesicles, astral microtubules and others (Eves et al., 2012). Underlined residues are important for vacuole inheritance, some of which were previously implicated in the transport of other cargoes, while others had not been tested (cyan). Among them, E1338, is predicted by AlphaFold to adopt a short helix (black) but it remains unresolved in the cryo-EM map and other myosin V tail structures. **(C)** Vacuole inheritance was tested for sites on Myo2 near the Vac17 density, Vac17(H). (n=3; ≥ 35 cells per n for each bud size). Significance was determined by unpaired, two-tailed, one-way ANOVA test with Dunnett’s multiple comparisons test. Error bars are SD. **** P < .001; ** P < .005; * P < .05; ns = not significant. **(D)** Representative blot of an *in vitro* pull-down assay of the Myo2 tail with Vac17(H) to assess the impact of these Myo2 mutations on the interaction. The wild-type and mutant of Strep II-Myo2 tail was co-expressed with MBP or MBP-Vac17(H) and immobilized on Strep-Tactin beads. **(E)** Binding efficiency from **(D)** was analyzed by normalizing to Vac17 and Myo2 input levels, respectively, to account for their variable expression levels. The intensity ratio of Vac17 pulled down with the Myo2 mutants was then compared as a fold change. Statistical analysis was performed using the Kruskal-Wallis test followed by Dunn’s multiple comparison test. **(F)** A suppressor mutation within Vac17(H), N60Y, rescues vacuole inheritance in the *myo2(E1333R, V1448A)* double mutant by binding the newly identified site. In a separate experiment, we found that a partial *vac17(F132Y)* mutant, located in the canonical Myo2-binding domain, when combined with *myo2(E1333R, V1448A)* at the Vac17(H) binding site, abolishes vacuole transport. This suggests that interactions at both sites are important for vacuole inheritance. **(G)** An AlphaFold model of the Vac17(H)-Myo2 interaction. This region of Vac17 forms an antiparallel 3-helix bundle. The predicted direct contact between Vac17 h2, residues 42-79, and Myo2 is supported by the observation that the *vac17(N60Y)* mutant in h2 rescues the vacuole inheritance defect of the *myo2(E1338R, V1448A)* mutant at this newly identified site. Note that Myo2(E1338) is absent from all resolved structures, and its precise location remains unknown. Green = Vac17(MBD) peptide; red = suppressor residues that may interact with Myo2; cyan = newly identified Myo2 residues; orange = Myo2 residue previously characterized in other cargo transport, also binds Vac17(H) and functions in vacuole inheritance.

Our cryo-EM analysis yielded a ∼7 Å reconstruction of the Vac17(H+MBD)-Myo2 complex, and revealed two distinct Vac17 densities surrounding Myo2 (**Fig. 5 B**). Given the low number of particles showing simultaneous occupancy at both sites, we infer that Vac17(H) is either more flexible or binds more weakly than Vac17(MBD). To further characterize this new interface, we docked predicted models of the Vac17(H+MBD)-Myo2 complex and found that Vac17(H) interacts with a region on Myo2 that was previously implicated in the transport of other cargoes, including secretory vesicles and peroxisomes.

We first tested Myo2 residues that likely contribute to this interface. Myo2(W1407) and Myo2(K1408) were previously shown to mediate the movement of astral microtubules and peroxisomes via Kar9 and Inp2, respectively (Eves et al., 2012; Fagarasanu et al., 2009). Notably, vacuole inheritance occurred in 63% and 57% small buds in the *myo2(W1407Y)* and *myo2(K1408A)* mutant cells, respectively (**Fig. 5, B and C**). These findings suggest that these Myo2 residues contribute to vacuole transport, in addition to their established roles in moving at least two other cargoes.

We also tested Myo2 residues that were not previously linked to cargo binding. *myo2(V1448A), myo2(R1449E), and myo2(I1458A)* mutants each resulted in a > 50% defect in vacuole inheritance. Additionally, the *myo2(E1338R*) mutation, which is located in an unresolved loop containing a regulatory phosphosite in myosin Va (**Fig. S3 F and Fig. 5 B;** (Karcher et al., 2001; Yoshizaki et al., 2007), impaired vacuole inheritance, with only 56% of vacuoles observed in the small buds.

To further test whether this newly identified region on Myo2 directly binds Vac17(H), we performed pull-down assays using recombinantly co-expressed Myo2 mutants with Vac17(H). Compared to wild-type Myo2 (**Fig. 5 D and E**), the *myo2(K1408A, R1449E)* double mutant and the *myo2(E1338R, K1408A, V1448A)* triple mutant tails exhibited progressively reduced binding with the Vac17(H) peptide. Together, these data establish a previously unrecognized Vac17(H)-binding site on Myo2.

To gain insights into how Vac17(H) engages this region, we analyzed multiple AlphaFold models, which revealed that Vac17(H) forms antiparallel 3-helix bundles in two distinct orientations relative to Myo2. We then mutated candidate Vac17 residues from all three helices—*vac17(E19K)*, *vac17(R30E, R33E)*, *vac17(N60Y)*, *vac17(S61E, V64E)*, and *vac17(E97R)*. Among these, only *vac17(N60Y)* rescued the vacuole inheritance defect of the *myo2(E1338R, V1448A)* mutant (**Fig. 5 F**). This suggests that h2, Vac17(42-79), directly interfaces with Myo2. Notably, all four original suppressor mutations in h2 substituted polar for hydrophobic residues. This suggests that these changes strengthen binding to the hydrophobic patch on Myo2—comprising Myo2(W1407), Myo2(V1448) and Myo2(I1458)—that is important for vacuole inheritance (**Fig. 5, B and G**). Similarly, the suppressor L94M in h3, is predicted to engage this same Myo2 surface.

The observation that suppressor mutations in Vac17(H) rescued a point mutation on the opposite side of Myo2, *myo2(N1304S)*, suggests that impaired interaction at one binding site, could be worsened by weakening the interaction at the other site. To test this, we examined whether a partial mutation in Vac17(MBD), *vac17(F132Y)*, would be further impaired by mutations in the newly identified Myo2 sites that interact with Vac17(H): *myo2(E133R, V1448A).* Notably, co-expression of these mutants resulted in a complete loss of vacuole inheritance (**Fig. 5, F and G**). These findings further support a model in which Vac17 engages Myo2 at two distinct sites during vacuole inheritance.

### Inp2 uses two independent sites on Myo2 for peroxisome transport

That Vac17 binds two different sites on the Myo2 tail raises the question of whether this feature is shared with other cargo adaptors. Indeed, the peroxisome adaptor Inp2 engages two separate sites on Myo2. *In vitro* studies revealed that the interaction of the Inp2(241-705) peptide with the *myo2(W1407F)* mutant tail was impaired, and in yeast the *myo2(W1407F)* mutant also resulted in a peroxisome inheritance defect (Eves et al., 2012; Fagarasanu et al., 2006; Fagarasanu et al., 2009). In contrast, a recent structural study revealed that the Inp2(531-543) peptide interacts at a distant site on Myo2, Myo2(F1264) and Myo2(F1275) (Tang et al., 2019; **Fig. 6 A**). These findings suggest that Myo2(F1264) and Myo2(F1275) may define a third cargo recognition site on Myo2, although this had not been tested *in vivo*.

**Figure 6.**
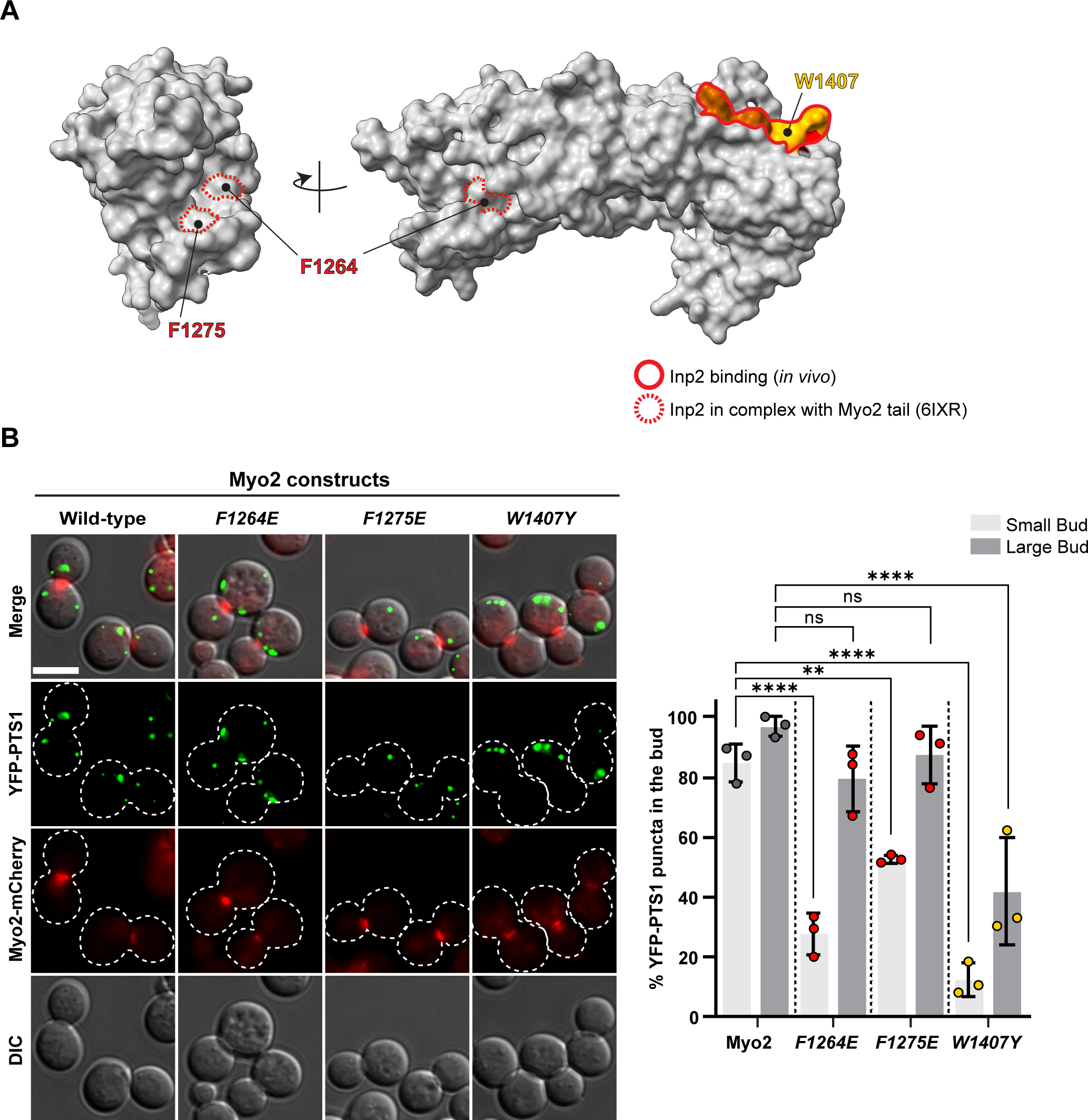
Peroxisome inheritance requires that the Inp2 adaptor binds to two distinct sites on Myo2. **(A)** Residues on the Myo2 tail identified as important for peroxisome inheritance *in vivo* (solid outline) are distinct from those observed in structural studies (dashed outline) of the Inp2 peptide(532-540) in complex with the Myo2 tail (PDB: 6IXR; Tang et al., 2019; Fagarasanu et al., 2006; Fagarasanu et al., 2009). **(B)** Myo2 residues implicated in Inp2 interaction, either through genetic studies, Myo2(W1407) or structural analysis, Myo2(F1264) and Myo2(F1275), were tested for their role in peroxisome inheritance. n=3; each n represents 40 ≥ cells per group. YFP-PTS1 encodes YFP fused with a peroxisome-targeting sequence. Scale bar = 5 μm.

To determine whether Inp2 spans Myo2 to engage two sites, we first tested *myo2(W1407Y),* and showed that there is a defect in peroxisome inheritance, with only 12% of small buds containing peroxisomes (**Fig. 6 B**). Moreover, in the *myo2(F1264E)* and *myo2(F1275E)* mutants, 28% and 53% of small buds contained peroxisomes. Note that in wild-type cells 82% of small buds contain peroxisomes (**Fig. 6 B**). These findings suggest that, similar to Vac17, Inp2 interacts with Myo2 at two distinct sites.

Cargo adaptors were previously thought to form a single interface with Myo2. However, our findings reveal that Vac17, as a single adaptor, forms extensive contacts and engages with at least two separate sites on Myo2. Importantly, Inp2 also binds to at least two sites on Myo2. This suggests a strategy where a single adaptor can form multiple distinct interfaces with Myo2. We propose that this handhold mechanism may govern the interactions of multiple cargo adaptors with myosin V.

## Discussion

Through a combination of genetics, live-cell imaging, biochemistry and structural biology, we discovered that Vac17 wraps around Myo2, engaging with two distinct interfaces on its tail. This “handhold” mechanism strengthens the motor-cargo association, and ensures efficient vacuole transport (**Fig. 7, A and B**). While Vac17(MBD) provides a high-affinity, essential interaction, it is insufficient for proper vacuole transport. This highlights the importance of multivalent interactions in at least some motor-cargo adaptor complexes. Importantly, Vac17(H) is required for optimal engagement with Myo2. Deletion of Vac17(H) results in a delay in vacuole movement (**Fig. 7, C and D**). We propose that the increased steady-state levels of the *vac17ΔH* mutant may eventually recruit sufficient Myo2 motors to move vacuoles into large buds. (**Fig. 7, E and F**).

**Figure 7.** Model for how Vac17 uses a handhold to stabilize its interaction with Myo2 and initiate vacuole movement into small buds. **(A)** Vacuole inheritance occurs concurrently with bud emergence or in small buds. The Myo2-Vac17-Vac8 complex localizes to a region of the vacuole near the bud site, and a portion of the vacuole is subsequently moved into the bud. **(B)** The proper timing of vacuole transport depends on stable interactions between Myo2 and Vac17. This is achieved by Vac17 engaging Myo2 at two distinct interfaces via its Vac1(H) and Vac17(MBD) regions. **(C, D)** In the *vac17ΔH* mutant, the initiation of vacuole movement is significantly delayed. This is likely due to a loss of stable interactions with Myo2, and highlights the critical role of the handhold mechanism in ensuring timely vacuole transport. **(E)** Vacuole inheritance eventually occurs in large buds of the *vac17ΔH* mutant. This is likely due to a defect in the turnover of Vac17, resulting in an abnormal accumulation of Vac17 on the vacuole membrane. **(F)** We speculate that this excess availability of Vac17 compensates for the partial defect in *vac17ΔH*, by promoting increased recruitment of Myo2 to the vacuole or by increasing the occupancy of Vac17(MBD) for Myo2.

Similarly, the peroxisome adaptor Inp2 also requires two Myo2-binding sites for efficient peroxisome transport. This provides another example of a single adaptor binding to multiple sites on Myo2. Previous structural studies suggested that each cargo adaptor engaged only one myosin V/Myo2 site (Pylypenko et al., 2013; Wei et al., 2013; Tang et al., 2019). However, these studies used adaptor peptides that were too short to encompass other potential binding sites. Our cryo-EM analysis enabled us to utilize a longer adaptor peptide, which revealed that Vac17 spans two Myo2 sites.

Despite successful crystallization of other adaptor peptides, Vac17 had remained structurally elusive. Cryo-EM provided molecular insights into Vac17-Myo2 interactions. While cryo-EM remains challenging for resolving high-resolution structures of small, asymmetric proteins, our reconstruction offers new insights by capturing the broader structural context of the interaction between Vac17 and Myo2.

Structural comparisons reveal that Vac17(MBD) uniquely binds Myo2 in an orientation opposite to other adaptors. Myo2-bound Vac17 and Mmr1, as well as myosin Va-bound Spir2 and Mlph utilize an equivalent hydrophobic groove, each stabilized by polar interactions on one side. This conserved binding pattern was not evident from sequence analysis alone.

Our analysis also reexamines phosphosites implicated in adaptor release (**Fig. S4 E**). Vac17(S131) and Mmr1(S414) align with each other and reside on the longest Myo2 tail helix. We hypothesize that these sites play a role in the dissociation of Vac17 or Mmr1 from Myo2, though likely in an indirect manner. Mmr1(pS414), which triggers ubiquitination and detachment from Myo2 during mitochondrial transport (Obara et al., 2022), faces away from Myo2 (Tang et al., 2019; Nayef et al., 2024), and thus, would not directly impact Mmr1 binding.

A second reconstruction, derived from 18% of particles, suggests that Vac17(H) is structurally flexible and may be stabilized by additional factors. This Vac17 density is further supported by AlphaFold models, which align with the hypothesis that Vac17(H) is positioned opposite and distal to the Vac17(MBD) site on Myo2. Vac17(H) likely interacts with Myo2 via hydrophobic contacts, specifically with Myo2(W1407), Myo2(V1448) and Myo2(I1458). Notably, many suppressor mutations identified in Vac17(H) involved changes to hydrophobic residues, suggesting that these mutations act by enhancing interaction with a hydrophobic surface on Myo2, including W1407, V1448 and I1458. In addition, Myo2(V1448) corresponds to the disease-associated *myosin Vb(L1746R)* residue (Aldrian et al., 2021), which may alter hydrophobic contacts with Rab11 (Pylypenko et al., 2013).

Another mutant, *myo2(E1338R)*, which impairs vacuole transport, is analogous to *myosin Vb(R1641C)*, a variant associated with cancer (Letellier et al., 2017). Both mutations are located in an unstructured region of the Myo2/myosin V tail, where a regulatory phosphosite in myosin Va(S1650) has been identified (Karcher et al., 2001; Yoshizaki et al., 2007). While Myo2(E1338) is distant from the hydrophobic patch, it likely contributes to Vac17(H) binding and plays a role in the transport of multiple cargoes. Together, these findings underscore the functional relevance of the newly characterized Myo2 residues.

Adaptor binding sites often overlap with key recognition interfaces on Myo2, suggesting that cargo recognition may be competitive. The observation that many adaptors either bind two sites—as shown here for the vacuole and peroxisome—or require two adaptors for a single organelle increases the potential for competition between adaptors. Specifically, Vac17(MBD) overlaps with the Mmr1-binding site, while Vac17(H) interacts near Ypt11. Since Mmr1 and Ypt11 cooperate to move mitochondria, (Chernyakov et al., 2013; Lewandowska et al., 2013), and live-cell imaging shows that vacuole inheritance generally precedes mitochondrial inheritance (Eves et al., 2012; Li et al., 2021), these overlapping adaptor interfaces likely facilitate temporal coordination of the transport of distinct organelles.

Our structural data suggest that a single Vac17 adaptor is unlikely to bridge the homodimeric tails of Myo2. If Myo2 adopts an autoinhibited conformation, similar to myosin Va (Niu et al., 2022), the distance between Vac17-binding sites would be too great for one Vac17 molecule to span both tails. Additionally, structural studies indicate that the C-terminus of Vac17 wraps around a single Vac8 (Kim et al., 2023). This supports a model in which a single Vac17 molecule tethers one Myo2 tail to Vac8.

In summary, our findings provide insights into the structural basis of motor-cargo interactions. Separate from the classical model in which multiple adaptors form independent contacts, our data support a handhold strategy in which a single adaptor engages multiple interactions. This novel feature of Vac17 interaction with Myo2 offers a new perspective on how myosin V-cargo interactions are achieved and thus regulated.

## Materials and Methods

### Cloning

Vac17 peptides, 1-109, 110-157, and 1-157, were PCR amplified from the Vac17 open reading frame, which was previously optimized for *E. coli* codon usage. Primer pairs included SalI and PstI restriction sites, and the corresponding PCR product was digested and ligated into an empty MBP-10xhis-TEV backbone. Strep II-Myo2(1150-1567) was generated using the Q5 SDM kit (NEB) to insert the epitope tag at the N-terminus of Myo2 open reading frame. Primers are listed in **Table S2**.

### Site-directed mutagenesis

Point mutations were introduced using the single-mutagenic primer method. Internal deletions or insertions were generated using the Q5 site-directed mutagenesis kit (NEB). Primer pairs were designed using NEBaseChanger.

### Strains and Plasmids

Strains used in this study are listed in **Table S3**. Plasmids used in this study are in **Table S4**. Yeast cultures were grown at 24°C at 200 rpm. Yeast extract-peptone-dextrose (YEPD) was prepared with the following reagents: 1% yeast extract, 2% peptone, 2% dextrose. Synthetic complete (SC) media lacking specific amino acid, and F-5OA plates for counter-selection were made as described (Adams et al., 1998).

### Protein expression and purification

MBP-10xhis-TEV-Vac17 constructs, and Strep II-Myo2 tail were co-expressed in BL21(DE3) cells. Primary cultures were grown overnight at 37°C in Luria Broth media at 180 rpm. The cultures were then backdiluted (100x) into Hyper Broth media. Once the OD_600_ reached ∼1.0, the shaker was chilled to 16°C, and cells were equilibrated for additional ∼0.5 hour, before induction with 0.1mM IPTG. After 18 hours of expression, cells were harvested by centrifugation at 4,000 rpm for 20 minutes, flash-frozen and stored at -80°C until protein purification. Pellets were weighed and resuspended in freshly prepared buffer containing 50mM phosphate, pH 7.6, 150 mM NaCl, 2 mM *β*-*Mercaptoethanol* (*β*ME), 1mM EDTA pH 8.0, 1 mM PMSF, EDTA-free protease inhibitor cocktail, and benzonase (25 U/ml). The pellets were processed through a high-pressure homogenizer (15 k psi; 2 passes). Cell lysates were cleared by ultracentrifugation at 40k rpm for 30 minutes at 4°C. The clarified lysates were incubated with pre-equilibrated amylose resin (NEB) for 3 hours at 4°C on a nutator. The resin was transferred to a 10 ml column and washed extensively, before adding TEV protease to cleave the MBP fusion tags, followed by 1.5 h of incubation at 16°C. Eluents were concentrated using a 10 kDa MWCO Amicon spin filter (MilliporeSigma). Analytes reaching ∼500 ul or 3 mg/ml were injected into a size-exclusion column (Superdex 200 Increase 10/300 GL) at 0.35 ml/min. Only fresh proteins from individual fractions were used for same-day cryo-EM grid preparation.

### *In vitro* binding assay

Flash-frozen pellets of BL21(DE3) cells co-expressing Strep II-Myo2 tail and MBP-Vac17(H) were resuspended in 750 μl of resuspension buffer (50mM Tris-HCl pH 8.0, 150mM NaCl, 5mM *β*ME, 1mM EDTA, 1mM PMSF, benzonase and protease inhibitor cocktail). The lysate was transferred to a 2 ml tube, and 250 μl of 0.1 mm zirconia beads were added for cell lysis using a Mini-BeadBeater-8 (agitate for 30 seconds, followed by cooling on ice, repeated six times). After lysis, the samples were centrifuged at 10,000 g for 5 minutes at 4°C, and the supernatant was transferred to a fresh 1.7 ml Eppendorf tube for a second round of centrifugation at 14,000 rpm for 10 minutes. The resulting soluble fraction was incubated with 100 μl of pre-equilibriated StreptactinXT 4Flow beads for 1 hour with gentle rotation at 4°C. The beads were then washed three times with 750 μl of wash buffer (50mM Tris-HCl pH 8.0, 400mM NaCl, 0.1% Tween, and 5mM *β*ME), with each wash step involving rotation for 3-5 minutes.

### Cryo-EM grid preparation and data acquisition

Freshly eluted protein fractions were individually sampled for cryo-EM grid preparation (∼0.15-0.3 mg/ml per fraction for 68 kDa sample). UltrAufoil R 1.2/1.3 on 300 mesh gold grids were glow discharged at 5 mA for 90 s and 10 s hold in a 0.26 mBar vacuum chamber. Initial structure determination attempts revealed a preferred orientation of the complex. To overcome this problem, *fluorinated octyl*-*maltoside* (FOM) [150 μM final] was added to the peak fraction, incubated on ice, and spun down to remove any precipitates, before sample vitrification. Grids were blotted with filter paper and plunge-frozen into ethane cooled with liquid nitrogen using a Vitrobot Mark IV (Thermo Fisher Scientific) set to 4°C, 100% humidity, with 0 s wait, blot 4 s, 20 force. Screening was performed overnight using a 200 keV Glacios or Arctica, using Leginon (Suloway et al., 2005). Data collection was done on a 300 keV Krios A equipped with a K3 Summit direct electron detector and a Gatan Imaging Filter with a 20 eV slit width in counting mode (1.085 *Å*/pixel) at nominal magnification of 81,000x. Beam-image shift was used for data collection.

### Image processing

For more details, see **Table S5** and **Fig. S5**. CryoSPARC (Punjani et al., 2017) v4 was used for preprocessing the dataset. After Patch Motion Correction and Patch CTF, exposures were manually curated. A random subset (10%) of micrographs was selected for blob picking to generate template particles for the curated dataset of 3,303 micrographs. After removing junk particles through three rounds of 2D classification, the remaining 618,699 particles were used for Ab-initio reconstruction and heterogeneous refinement, including junk volume. The highest resolution map, comprising 200,770 particles, underwent 2D classification, and several rounds of heterogeneous refinements. The resulting 144,724 particles were selected for non-uniform refinement to obtain a final resolution of 5.75 Å. To obtain the fuzzy density structure, the original density for Myo2 and Vac17(MBD) from 200,770 particles was subtracted from particle images. Using ChimeraX’s molmap function, a composite of AlphaFold models for Vac17(1-09) was used as a template to generate a 15 Å Vac17 map, which was positioned near the fuzzy density in the un-subtracted reconstruction. The map was converted into a mask in CryoSPARC and further expanded by dilating and applying soft padding, for alignment-free focused 3D classification. For the highest resolution model, representing 35,957 particles, un-subtracted particle images were reintroduced for homogenous reconstruction. Following non-uniform refinement, the final map resolved to 7.56 Å. A test *ab-initio* reconstruction with 35,957 particles from the refinement job recapitulated the final map.

### Vacuole inheritance assay

Primary overnight cultures grown to 0.2-0.35 OD_600_/ml, were used for vacuole labeling. Cells were resuspended in 250 μl of fresh selective media into 1.7 ml Eppendorf tubes, and 3 μl of FM 4-64 dye (in DMSO; SynaptoRed C2, Biotium, Inc.) was added to a final concentration of 24 nM. The tubes were wrapped in aluminum foil and incubated for 40 minutes at 24°C on a shaker at 200 rpm. After incubation, cells were washed twice with fresh media, resuspended in 1ml, and backdiluted into 4 ml of fresh media (1:5). Cells were incubated for at least one doubling time (3 h), before imaging.

### Alpha-factor synchronization for time-lapse imaging

Primary overnight cultures grown to 0.2-0.3 OD_600_/ml (50ml) were synchronized in G1 phase by incubating with fresh aliquots of 4 μM of alpha-factor (Zymo Research) for 3 h. Cells were washed three times with fresh media, resuspended several times to break up clumps, and transferred into a microfluidics plate (200 μl total volume), which was continuously perfused with new media. Cells were monitored for one doubling cycle before imaging to avoid an alpha-factor induced accumulation of Vac17 on vacuole transport. 10-15 OD’s of cells were loaded into the CellASIC ONIX2 microfluidic plate (Millipore) for time-lapse imaging. Cells were viewed with DeltaVision microscope using a 60x objective and 1.6x auxiliary magnification (0.06788 μm/pixel).

### Epifluorescence Microscopy/Image Processing

Yeast cells were imaged using a DeltaVision Restoration system (Applied Precision) using an inverted epifluorescence microscope (IX-71; Olympus), equipped with a 100x objective and CCD camera (Cool-SNAP HQ; Photometrics). For PTS1 and Myo2 puncta visualization (**Fig. 6**), imaging was performed on a Leica DMi8 microscope with a 100x objective and Leica Thunder imager. Single-z sections were taken for all imaging modalities. Images were processed and analyzed using FIJI (Schindelin et al., 2012).

### Yeast Whole Cell Lysate Extraction

Cells at 0.4-0.6 OD_600_/ml (50 ml total) were harvested by centrifugation at 3,000 g for 3 minutes and flash frozen. The pellet was resuspended in 1 ml prechilled lysis buffer (0.2 M NaCl, 7.5% *β*ME) and incubated on ice for 10 minutes. Then, 100 μl of prechilled trichloroacetic acid (TCA) was added and left for an additional 10 minutes. The lysate was centrifuged at 13k rpm for 5 minutes at room temperature. The supernatant was discarded, and the pellet was air-dried for ∼15 minutes. The pellets were resuspended in 100 μl of 2X sample buffer (4% SDS, 0.1 M Tris-HCl pH 6.8, 20% glycerol, 0.01% Bromophenol Blue, 5% *β*ME) and 1M Tris base (pH 11) was added incrementally (20 μl) to neutralize the pH of the precipitates. Samples were boiled at 75°C for 10 minutes, spun down, and loaded onto SDS-PAGE gels, which were run at 150V for 1.5 h.

### Protein Electrophoresis and Western Blot

For direct protein detection, SDS-PAGE gels were incubated with Imperial Protein Stain (Thermo Scientific) and destained with ddH2O. For immunoblot analysis, gels were transferred onto 0.45 μm nitrocellulose membranes by wet transfer at 100V for 3 h. Membranes were blocked at room temperature for 45 minutes with 5% milk in 1x TBST. Primary antibodies used were rabbit ɑ-Vac17 (1:3000), goat ɑ-Myo2 (1:3000), mouse ɑ-GFP (1:1000; Roche) and mouse ɑ-Pgk1 (1:10000; Invitrogen), and were incubated for 3 h at room temperature or 16 h in the cold room. Membranes were washed three times with 1x TBST at 10-minute intervals, followed by incubation with secondary antibodies (1:5000-1:10000) for 1 hour in fresh milk at room temperature. Blots were washed three times in TBST. Proteins were detected with standard ECL (Cytiva), except for GFP and Vac17 signal from yeast lysates, which were detected using ECL prime. Blots were developed with the ChemiDoc instrument and quantified with the BioRad Image Lab Software.

## Data Availability Statement

All data are available from the corresponding authors upon request.

## Acknowledgements

We thank Drs. Shyamal Mosalaganti, Amir Khan, Ming Li and members of the Weisman and Cianfrocco labs for their insightful comments. We thank Ming Li for providing access to the CellASIC ONIX2 microfluidic system for the live-cell imaging experiments. The research reported in this publication was supported by the University of Michigan Cryo-EM Facility (U-M Cryo-EM). U-M Cryo-EM is grateful for support from the U-M Life Sciences Institute and the U-M Biosciences Initiative. We thank Alim Habib for generating the 3xEnvy construct, Dr. Yui Jin for the pRS413-mCherry-Myo2 plasmid, and Dr. Amir Khan for the pRSF-Duet-1 plasmid, and codon optimized Vac17 for expression in *E. coli*. We thank Dr. Charles G. Krasnow for suggesting the term “handhold” mechanism for describing our model. This work was supported by the National Institutes of Health grant R01-GM062261 to L.S.W and S10OD020011. H.J.H was supported in part by the University of Michigan Rackham Predoctoral Fellowship.

## References

Adams, A., C. Kaiser, and Cold Spring Harbor Laboratory. 1998. Methods in yeast genetics : a Cold Spring Harbor Laboratory course manual. Cold Spring Harbor Laboratory Press, Plainview, N.Y. xiv, 177 p. pp.

Aldrian, D., G.F. Vogel, T.K. Frey, H. Ayyildiz Civan, A.U. Aksu, Y. Avitzur, E. Ramos Boluda, M. Cakir, A.M. Demir, C. Deppisch, H.C. Duba, G. Duker, P. Gerner, J. Hertecant, J. Hornova, S. Kathemann, J. Koeglmeier, A. Koutroumpa, R. Lanzersdorfer, R. Lev-Tzion, R. Lima, S. Mansour, M. Meissl, J. Melek, M. Miqdady, J.H. Montoya, C. Posovszky, Y. Rachman, T. Siahanidou, M. Tabbers, H.H. Uhlig, S. Unal, S. Wirth, F.M. Ruemmele, M.W. Hess, L.A. Huber, T. Muller, E. Sturm, and A.R. Janecke. 2021. Congenital Diarrhea and Cholestatic Liver Disease: Phenotypic Spectrum Associated with MYO5B Mutations. J Clin Med. 10.

Beach, D.L., J. Thibodeaux, P. Maddox, E. Yeh, and K. Bloom. 2000. The role of the proteins Kar9 and Myo2 in orienting the mitotic spindle of budding yeast. Curr Biol. 10:1497–1506.

Beningo, K.A., S.H. Lillie, and S.S. Brown. 2000. The yeast kinesin-related protein Smy1p exerts its effects on the class V myosin Myo2p via a physical interaction. Mol Biol Cell. 11:691–702.

Catlett, N.L., J.E. Duex, F. Tang, and L.S. Weisman. 2000. Two distinct regions in a yeast myosin-V tail domain are required for the movement of different cargoes. J Cell Biol. 150:513–526.

Catlett, N.L., and L.S. Weisman. 1998. The terminal tail region of a yeast myosin-V mediates its attachment to vacuole membranes and sites of polarized growth. Proc Natl Acad Sci U S A. 95:14799–14804.

Chernyakov, I., F. Santiago-Tirado, and A. Bretscher. 2013. Active segregation of yeast mitochondria by Myo2 is essential and mediated by Mmr1 and Ypt11. Curr Biol. 23:1818–1824.

Cianfrocco, M.A., Wong-Barnum, M., Youn, C., Wagner, R. and Leschziner, A.. 2017. COSMIC2: A Science Gateway for Cryo-Electron Microscopy Structure Determination. In PEARC17: Practice and Experience in Advanced Research Computing 2017. Association for Computing Machinery, New Orleans, LA. 5.

Donovan, K.W., and A. Bretscher. 2015. Head-to-tail regulation is critical for the in vivo function of myosin V. J Cell Biol. 209:359–365.

Dunkler, A., M. Leda, J.M. Kromer, J. Neller, T. Gronemeyer, A.B. Goryachev, and N. Johnsson. 2021. Type V myosin focuses the polarisome and shapes the tip of yeast cells. J Cell Biol. 220.

Engevik, A.C., I. Kaji, M.M. Postema, J.J. Faust, A.R. Meyer, J.A. Williams, G.N. Fitz, M.J. Tyska, J.M. Wilson, and J.R. Goldenring. 2019. Loss of myosin Vb promotes apical bulk endocytosis in neonatal enterocytes. J Cell Biol. 218:3647–3662.

Evans, R., M. O’Neill, A. Pritzel, N. Antropova, A. Senior, T. Green, A. Žídek, R. Bates, S. Blackwell, J. Yim, O. Ronneberger, S. Bodenstein, M. Zielinski, A. Bridgland, A. Potapenko, A. Cowie, K. Tunyasuvunakool, R. Jain, E. Clancy, P. Kohli, J. Jumper, and D. Hassabis. 2022. Protein complex prediction with AlphaFold-Multimer. bioRxiv:2021.2010.2004.463034.

Eves, P.T., Y. Jin, M. Brunner, and L.S. Weisman. 2012. Overlap of cargo binding sites on myosin V coordinates the inheritance of diverse cargoes. J Cell Biol. 198:69–85.

Fagarasanu, A., M. Fagarasanu, G.A. Eitzen, J.D. Aitchison, and R.A. Rachubinski. 2006. The peroxisomal membrane protein Inp2p is the peroxisome-specific receptor for the myosin V motor Myo2p of Saccharomyces cerevisiae. Dev Cell. 10:587–600.

Fagarasanu, A., F.D. Mast, B. Knoblach, Y. Jin, M.J. Brunner, M.R. Logan, J.N. Glover, G.A. Eitzen, J.D. Aitchison, L.S. Weisman, and R.A. Rachubinski. 2009. Myosin-driven peroxisome partitioning in S. cerevisiae. J Cell Biol. 186:541–554.

Fortsch, J., E. Hummel, M. Krist, and B. Westermann. 2011. The myosin-related motor protein Myo2 is an essential mediator of bud-directed mitochondrial movement in yeast. J Cell Biol. 194:473–488.

Hammer, J.A., 3rd, and J.R. Sellers. 2011. Walking to work: roles for class V myosins as cargo transporters. Nat Rev Mol Cell Biol. 13:13-26.

Hill, K.L., N.L. Catlett, and L.S. Weisman. 1996. Actin and myosin function in directed vacuole movement during cell division in Saccharomyces cerevisiae. J Cell Biol. 135:1535–1549.

Holthenrich, A., J. Terglane, J. Nass, M. Mietkowska, E. Kerkhoff, and V. Gerke. 2022. Spire1 and Myosin Vc promote Ca(2+)-evoked externalization of von Willebrand factor in endothelial cells. Cell Mol Life Sci. 79:96.

Ishikawa, K., N.L. Catlett, J.L. Novak, F. Tang, J.J. Nau, and L.S. Weisman. 2003. Identification of an organelle-specific myosin V receptor. J Cell Biol. 160:887–897.

Itoh, T., E.A. Toh, and Y. Matsui. 2004. Mmr1p is a mitochondrial factor for Myo2p-dependent inheritance of mitochondria in the budding yeast. EMBO J. 23:2520–2530.

Itoh, T., A. Watabe, E.A. Toh, and Y. Matsui. 2002. Complex formation with Ypt11p, a rab-type small GTPase, is essential to facilitate the function of Myo2p, a class V myosin, in mitochondrial distribution in Saccharomyces cerevisiae. Mol Cell Biol. 22:7744–7757.

Jin, Y., A. Sultana, P. Gandhi, E. Franklin, S. Hamamoto, A.R. Khan, M. Munson, R. Schekman, and L.S. Weisman. 2011. Myosin V transports secretory vesicles via a Rab GTPase cascade and interaction with the exocyst complex. Dev Cell. 21:1156–1170.

Jin, Y., and L.S. Weisman. 2015. The vacuole/lysosome is required for cell-cycle progression. Elife. 4.

Jumper, J., R. Evans, A. Pritzel, T. Green, M. Figurnov, O. Ronneberger, K. Tunyasuvunakool, R. Bates, A. Zidek, A. Potapenko, A. Bridgland, C. Meyer, S.A.A. Kohl, A.J. Ballard, A. Cowie, B. Romera-Paredes, S. Nikolov, R. Jain, J. Adler, T. Back, S. Petersen, D. Reiman, E. Clancy, M. Zielinski, M. Steinegger, M. Pacholska, T. Berghammer, S. Bodenstein, D. Silver, O. Vinyals, A.W. Senior, K. Kavukcuoglu, P. Kohli, and D. Hassabis. 2021. Highly accurate protein structure prediction with AlphaFold. Nature. 596:583–589.

Karcher, R.L., J.T. Roland, F. Zappacosta, M.J. Huddleston, R.S. Annan, S.A. Carr, and V.I. Gelfand. 2001. Cell cycle regulation of myosin-V by calcium/calmodulin-dependent protein kinase II. Science. 293:1317–1320.

Kim, H., J. Park, H. Kim, N. Ko, J. Park, E. Jang, S.Y. Yoon, J.A.R. Diaz, C. Lee, and Y. Jun. 2023. Structures of Vac8-containing protein complexes reveal the underlying mechanism by which Vac8 regulates multiple cellular processes. Proc Natl Acad Sci U S A. 120:e2211501120.

Lapierre, L.A., R. Kumar, C.M. Hales, J. Navarre, S.G. Bhartur, J.O. Burnette, D.W. Provance, Jr., J.A. Mercer, M. Bahler, and J.R. Goldenring. 2001. Myosin vb is associated with plasma membrane recycling systems. Mol Biol Cell. 12:1843–1857.

Letellier, E., M. Schmitz, A. Ginolhac, F. Rodriguez, P. Ullmann, K. Qureshi-Baig, S. Frasquilho, L. Antunes, and S. Haan. 2017. Loss of Myosin Vb in colorectal cancer is a strong prognostic factor for disease recurrence. Br J Cancer. 117:1689–1701.

Lewandowska, A., J. Macfarlane, and J.M. Shaw. 2013. Mitochondrial association, protein phosphorylation, and degradation regulate the availability of the active Rab GTPase Ypt11 for mitochondrial inheritance. Mol Biol Cell. 24:1185–1195.

Li, K.W., M.S. Lu, Y. Iwamoto, D.G. Drubin, and R.T.A. Pedersen. 2021. A preferred sequence for organelle inheritance during polarized cell growth. J Cell Sci. 134.

Lipatova, Z., A.A. Tokarev, Y. Jin, J. Mulholland, L.S. Weisman, and N. Segev. 2008. Direct interaction between a myosin V motor and the Rab GTPases Ypt31/32 is required for polarized secretion. Mol Biol Cell. 19:4177–4187.

Liu, J., D.W. Taylor, E.B. Krementsova, K.M. Trybus, and K.A. Taylor. 2006. Three-dimensional structure of the myosin V inhibited state by cryoelectron tomography. Nature. 442:208–211.

Liu, Y., L. Li, C. Yu, F. Zeng, F. Niu, and Z. Wei. 2022. Cargo Recognition Mechanisms of Yeast Myo2 Revealed by AlphaFold2-Powered Protein Complex Prediction. Biomolecules. 12.

Mirdita, M., K. Schutze, Y. Moriwaki, L. Heo, S. Ovchinnikov, and M. Steinegger. 2022. ColabFold: making protein folding accessible to all. Nat Methods. 19:679–682.

Nalavadi, V.C., L.E. Griffin, P. Picard-Fraser, A.M. Swanson, T. Takumi, and G.J. Bassell. 2012. Regulation of zipcode binding protein 1 transport dynamics in axons by myosin Va. J Neurosci. 32:15133–15141.

Nayef, N., L. Ekal, E.H. Hettema, and K.R. Ayscough. 2024. Insights into the regulation of the mitochondrial inheritance and trafficking adaptor protein Mmr1 in Saccharomyces cerevisiae. Kinases and Phosphatases. 2:190–208.

Niu, F., Y. Liu, K. Sun, S. Xu, J. Dong, C. Yu, K. Yan, and Z. Wei. 2022. Autoinhibition and activation mechanisms revealed by the triangular-shaped structure of myosin Va. Sci Adv. 8:eadd4187.

Obara, K., T. Yoshikawa, R. Yamaguchi, K. Kuwata, K. Nakatsukasa, K. Nishimura, and T. Kamura. 2022. Proteolysis of adaptor protein Mmr1 during budding is necessary for mitochondrial homeostasis in Saccharomyces cerevisiae. Nat Commun. 13:2005.

Otzen, M., R. Rucktaschel, S. Thoms, K. Emmrich, A.M. Krikken, R. Erdmann, and I.J. van der Klei. 2012. Pex19p contributes to peroxisome inheritance in the association of peroxisomes to Myo2p. Traffic. 13:947–959.

Pashkova, N., N.L. Catlett, J.L. Novak, and L.S. Weisman. 2005. A point mutation in the cargo-binding domain of myosin V affects its interaction with multiple cargoes. Eukaryot Cell. 4:787–798.

Pashkova, N., Y. Jin, S. Ramaswamy, and L.S. Weisman. 2006. Structural basis for myosin V discrimination between distinct cargoes. EMBO J. 25:693–700.

Peng, Y., and L.S. Weisman. 2008. The cyclin-dependent kinase Cdk1 directly regulates vacuole inheritance. Dev Cell. 15:478–485.

Punjani, A., J.L. Rubinstein, D.J. Fleet, and M.A. Brubaker. 2017. cryoSPARC: algorithms for rapid unsupervised cryo-EM structure determination. Nat Methods. 14:290–296.

Pylypenko, O., W. Attanda, C. Gauquelin, M. Lahmani, D. Coulibaly, B. Baron, S. Hoos, M.A. Titus, P. England, and A.M. Houdusse. 2013. Structural basis of myosin V Rab GTPase-dependent cargo recognition. Proc Natl Acad Sci U S A. 110:20443–20448.

Pylypenko, O., T. Welz, J. Tittel, M. Kollmar, F. Chardon, G. Malherbe, S. Weiss, C.I. Michel, A. Samol-Wolf, A.T. Grasskamp, A. Hume, B. Goud, B. Baron, P. England, M.A. Titus, P. Schwille, T. Weidemann, A. Houdusse, and E. Kerkhoff. 2016. Coordinated recruitment of Spir actin nucleators and myosin V motors to Rab11 vesicle membranes. Elife. 5.

Rogers, S.L., R.L. Karcher, J.T. Roland, A.A. Minin, W. Steffen, and V.I. Gelfand. 1999. Regulation of melanosome movement in the cell cycle by reversible association with myosin V. J Cell Biol. 146:1265–1276.

Roland, J.T., D.M. Bryant, A. Datta, A. Itzen, K.E. Mostov, and J.R. Goldenring. 2011. Rab GTPase-Myo5B complexes control membrane recycling and epithelial polarization. Proc Natl Acad Sci U S A. 108:2789–2794.

Schindelin, J., I. Arganda-Carreras, E. Frise, V. Kaynig, M. Longair, T. Pietzsch, S. Preibisch, C. Rueden, S. Saalfeld, B. Schmid, J.Y. Tinevez, D.J. White, V. Hartenstein, K. Eliceiri, P. Tomancak, and A. Cardona. 2012. Fiji: an open-source platform for biological-image analysis. Nat Methods. 9:676-682.

Schott, D., J. Ho, D. Pruyne, and A. Bretscher. 1999. The COOH-terminal domain of Myo2p, a yeast myosin V, has a direct role in secretory vesicle targeting. J Cell Biol. 147:791–808.

Spellman, P.T., G. Sherlock, M.Q. Zhang, V.R. Iyer, K. Anders, M.B. Eisen, P.O. Brown, D. Botstein, and B. Futcher. 1998. Comprehensive identification of cell cycle-regulated genes of the yeast Saccharomyces cerevisiae by microarray hybridization. Mol Biol Cell. 9:3273–3297.

Suloway, C., J. Pulokas, D. Fellmann, A. Cheng, F. Guerra, J. Quispe, S. Stagg, C.S. Potter, and B. Carragher. 2005. Automated molecular microscopy: the new Leginon system. J Struct Biol. 151:41–60.

Tang, F., E.J. Kauffman, J.L. Novak, J.J. Nau, N.L. Catlett, and L.S. Weisman. 2003. Regulated degradation of a class V myosin receptor directs movement of the yeast vacuole. Nature. 422:87–92.

Tang, K., Y. Li, C. Yu, and Z. Wei. 2019. Structural mechanism for versatile cargo recognition by the yeast class V myosin Myo2. J Biol Chem. 294:5896–5906.

Thirumurugan, K., T. Sakamoto, J.A. Hammer, 3rd, J.R. Sellers, and P.J. Knight. 2006. The cargo-binding domain regulates structure and activity of myosin 5. Nature. 442:212-215.

Trybus, K.M. 2008. Myosin V from head to tail. Cell Mol Life Sci. 65:1378–1389.

Varadi, A., T. Tsuboi, and G.A. Rutter. 2005. Myosin Va transports dense core secretory vesicles in pancreatic MIN6 beta-cells. Mol Biol Cell. 16:2670–2680.

Vida, T.A., and S.D. Emr. 1995. A new vital stain for visualizing vacuolar membrane dynamics and endocytosis in yeast. J Cell Biol. 128:779–792.

Wagner, W., S.D. Brenowitz, and J.A. Hammer, 3rd. 2011. Myosin-Va transports the endoplasmic reticulum into the dendritic spines of Purkinje neurons. Nat Cell Biol. 13:40-48.

Wang, Y.X., N.L. Catlett, and L.S. Weisman. 1998. Vac8p, a vacuolar protein with armadillo repeats, functions in both vacuole inheritance and protein targeting from the cytoplasm to vacuole. J Cell Biol. 140:1063–1074.

Wei, Z., X. Liu, C. Yu, and M. Zhang. 2013. Structural basis of cargo recognitions for class V myosins. Proc Natl Acad Sci U S A. 110:11314–11319.

Weisman, L.S. 2006. Organelles on the move: insights from yeast vacuole inheritance. Nat Rev Mol Cell Biol. 7:243–252.

Wong, S., N.L. Hepowit, S.A. Port, R.G. Yau, Y. Peng, N. Azad, A. Habib, N. Harpaz, M. Schuldiner, F.M. Hughson, J.A. MacGurn, and L.S. Weisman. 2020. Cargo Release from Myosin V Requires the Convergence of Parallel Pathways that Phosphorylate and Ubiquitylate the Cargo Adaptor. Curr Biol. 30:4399–4412 e4397.

Wu, X., B. Bowers, Q. Wei, B. Kocher, and J.A. Hammer, 3rd. 1997. Myosin V associates with melanosomes in mouse melanocytes: evidence that myosin V is an organelle motor. J Cell Sci. 110 (Pt 7):847-859.

Yau, R.G., Y. Peng, R.R. Valiathan, S.R. Birkeland, T.E. Wilson, and L.S. Weisman. 2014. Release from myosin V via regulated recruitment of an E3 ubiquitin ligase controls organelle localization. Dev Cell. 28:520–533.

Yau, R.G., S. Wong, and L.S. Weisman. 2017. Spatial regulation of organelle release from myosin V transport by p21-activated kinases. J Cell Biol. 216:1557–1566.

Yin, H., D. Pruyne, T.C. Huffaker, and A. Bretscher. 2000. Myosin V orientates the mitotic spindle in yeast. Nature. 406:1013–1015.

Yoshizaki, T., T. Imamura, J.L. Babendure, J.C. Lu, N. Sonoda, and J.M. Olefsky. 2007. Myosin 5a is an insulin-stimulated Akt2 (protein kinase Bbeta) substrate modulating GLUT4 vesicle translocation. Mol Cell Biol. 27:5172–5183.

