## Supplemental Tables and Figures for "Cargo adaptors use a handhold mechanism to engage with myosin V for organelle transport"

**Supplement**

**Table S1** shows Vac17 suppressor mutant candidates identified from the PCR screen. The percentage of cells containing an inherited vacuole in the bud was measured. For each group, between 30-200 cells were analyzed, with data collected from at least two biological replicates. For more specific details about the screen, see (Ishikawa et al., 2003). **Table S2** lists the primers used for this study. **Table S3** lists strains and **Table S4** lists plasmids used in this study. **Table S5** contains information regarding cryo-EM data collection, refinement and statistics. **Fig. S1** shows that the *vac17ΔH* mutant is defective in its localization to sites of polarized growth, but not with Vac8, and that Vac17(H) binds directly to the Myo2 tail *in vitro*. **Fig. S2** shows that the expression of Vac17 in the *vac17ΔH* mutant is elevated, and that the vacuole inheritance in small buds is delayed in the mutant. **Fig. S3** shows an overview of the cryo-EM specimen's purification, data collection. refinement and validation statistics. **Fig. S4** shows that Vac17 and Mmr1 bind overlapping sites on Myo2. **Fig. S5** shows cryo-EM processing workflow and validation statistics for the composite map of Vac17 bound to Myo2 at two sites.

**Figure Legends**

**Supplementary Figure 1. the *vac17ΔH* mutation specifically impairs its interaction with Myo2.**

**(A)** In wild-type cells, Vac17 accumulates on the vacuole surface closest to the bud cortex, while Myo2 localizes to either the bud tip or the mother-bud neck. Upon vacuole arrival at the bud tip, Vac17 undergoes degradation. In the *myo2(D1297N)* mutant, which is defective in vacuole inheritance, Vac17 is distributed around the vacuole, whereas *myo2(D1297N)* maintains its normal localization at sites of polarized growth. The *vac17ΔH* and *myo2(D1297N)* double mutant does not further alter the localization pattern of Vac17 or Myo2 compared to the *myo2(D1297N)* mutant. Scale bars = 5 µm.

**(B)** *vac17ΔH* still colocalizes with Vac8-RFP (pseudo-colored yellow), indicating that its binding to Vac8 is not impaired.

**(C)** In the absence of Vac8, Vac17 localizes presumably with Myo2, at the bud tip or the mother-bud neck, but *vac17ΔH* is cytosolic. Scale bars = 5 µm.

**(D)** Deletion of the canonical Vac17(MBD) abolishes vacuole inheritance.

**(E)** (Top) SDS-PAGE analysis of apo Myo2 tail purification. The peak elutes at 15.5 -16 ml. (Middle) Analysis of the Myo2 tail+Vac17(MBD) complex, showing an expected peak between 15.5 - 16 ml. There is also a higher molecular weight complex with both Vac17 and Myo2 at 13.5 - 14 ml. This latter peak was not analyzed further. (Bottom) Immunoblot analysis of the gel filtration experiment from (**Fig. 1 E**; top panel, lighter blue trace) shows the Myo2 tail+Vac17(H) complex at 14.5 - 15 ml. The membrane used to detect Myo2 was stripped and re-probed for MBP. The arrowheads correspond with the same arrowheads shown in (**Fig. 1 E)**: Gray = apo Myo2 tail; orange = free MBP; blue = Myo2 tail+Vac17(H); green = Myo2 tail+Vac17(MBD).

**Supplementary Figure 2. Elevated levels of Vac17 in the *vac17ΔH* mutant may eventually suppress the vacuole inheritance defect.**

**(A-C)** Buds that ultimately received vacuoles were chosen from (**Fig. 2, A and C)** and analyzed retrospectively.

**(A, B)** Vacuole inheritance over time following bud emergence (n=24 cells for wild-type; n=7 for *vac17ΔH* mutants). Time 0 represents cells where vacuole inheritance occurred concomitantly with bud emergence. After time 0, all cells that achieved vacuole inheritance at any time were included.

**(C)** Analysis of the timing of vacuole inheritance in large buds of *vac17ΔH* mutant cells that lacked a vacuole at the start of imaging (n=13). Large buds inherited vacuoles at times ranging from 20 seconds to 40 minutes.

**(D)** Western blot analysis of lysates from cells expressing Vac17 from a CEN (low copy) plasmid, compared to vector alone (n=3). Cells were harvested during mid-log phase ($OD_{600}$ 0.4-0.6). The vector control did not contain detectable bands. This confirms that the doublet bands in Vac17-3xEnvy and *vac17ΔH-3xEnvy* are Vac17 protein. Both bands were included in the quantification. Deletion of the PEST motif, which is essential for Vac17 degradation, results in elevated Vac17 levels.

**Supplementary Figure 3. Purification and cryo-EM analysis of the Vac17-Myo2 complex.**

**(A)** Example of preparative gel filtration showing co-expression of recombinant Vac17(H+MBD) and Myo2 tail. The black arrowhead indicates the complex formation for Vac17(H+MBD) and Myo2, which was sampled for data collection (top). The orange arrowhead marks free MBP, and the gray arrowhead indicates apo Myo2 tail. SDS-PAGE analysis of fractions from volumes 10-18 ml is shown.

**(B)** Example micrograph with selected particles. Scale bar = 100 nm.

**(C)** 2D class averages of particles processed using CryoSPARC. Scale bar = 130 Å.

**(D)** Local resolution range of the reconstruction.

**(E)** Model statistics (top) and Euler distribution of particles.

**(F)** Comparison of the Myo2 tail (2F6H; yellow) with the AlphaFold model, trimmed to fit the cryo-EM reconstruction (gray). Black circles indicate regions resolved in the cryo-EM map. Red circle highlights a predicted region lacking empirical data.

**Supplementary Figure 4. Mmr1 and Vac17(MBD) bind a shared site on Myo2 and are anti-parallel to each other.**

**(A)** AlphaFold model docked within the cryo-EM map of Vac17(MBD; 127-147) bound to Myo2 tail. Vac17-binding region (blue), Mmr1-binding region (red) and overlapping region of Vac17 and Mmr1 on Myo2 (pink) (Eves et al., 2012; this study).

**(B)** Vac17(MBD) extends along a hydrophobic groove (gold) on the Myo2 tail, which contains multiple surface residues critical for vacuole inheritance.

**(C)** Vac17(L137) makes hydrophobic interactions with Myo2(L1301 and L1229).

**(D)** An Mmr1 peptide(408-425) bound to the Myo2 tail (PDB: 6IXP) was docked into the EM reconstruction. The Mmr1 peptide partially fits into the density of Vac17(MBD), consistent with a previous study showing that Vac17 and Mmr1 peptides compete for access to Myo2 *in vitro* (Eves et al., 2012).

**(E)** Both the Vac17 and Mmr1 adaptor proteins contain serine phosphosites (depicted as space-filing model) that are important for dissociation from Myo2 (Wong et al., 2020; Obara et al., 2022).

**Supplementary Figure 5. Vac17-Myo2 cryo-EM data processing workflow.**

**Table S1 Vac17 suppressor screen**

| % Vacuole inheritance | | |
|---|---|---|
| VAC17 Suppressor candidates | | |
| | wild-type Myo2 | *myo2-N1304S* |
| I15V | 96 | 48 |
| I28R | 94 | 41 |
| N53Y | 95 | 50 |
| Y55H | 95 | 51 |
| S57F | 93 | 60 |
| N60Y | 94 | 58 |
| N60I | 98 | 49 |
| L96M | 97 | 57 |
| T110P | 93 | 66 |
| R126S | 95 | 51 |
| I140V | 95 | 52 |

**Table S2 Select primers used in this study**

| | | |
|---|---|---|
| *In vitro* **studies** | | |
| LHP211 | F SalI - vac17(1-109) - PstI | tcagacGTCGACATGGCTACGCAAGCCCTGGAAGAC |
| LHP212 | R SalI - vac17(1-109) - PstI | tcagacCTGCAGTTATGCCAGTTCATTCAGGCGCAGGGT |
| LHP216 | F SalI - vac17(110-157) - PstI | tcagacGTCGACACGGTGCCGAATGAAGCTAGTAACG |
| LHP217 | R SalI - vac17(157) - PstI | tcagacCTGCAGTTAGGATTTTTCGGCGGTTTACTCGGGG |
| LHP364 | myo2 tail-V1448A | GCTACTGCAAGcCCGTAAGTATAC |
| LHP377 | myo2 tail-R1449E | GCTACTGCAAGTCgaaAAGTATACTATCGAAGAC |
| LHP378 | myo2 tail-K1408A | TAATTTCTTGTCGTGGgcAAGGGGTCTT |
| LHP380 | myo2 tail-E1338R | CGCGGGAcgAACCAGCGGGTTTTT |
| *In vivo* **studies** | | |
| LHP319 | myo2-W1407Y | CGTAATTTCTTGTCGTacAAAAGGGGTCTTC |
| LHP285 | myo2-E1222K | GGCTGACCAAGCAAAGTaAAAGCTTTCTTGCCC |
| LHP275 | myo2-V1448A | CCGCTAAGCTACTGCAAGcCCGTAAGTATACTATCG |
| LHP277 | myo2-R1449E | GACCGCTAAGCTACTGCAAGTCgaaAAGTATACTATCGAAGAC |
| LHP280 | myo2-I1462R | CGAAGACATTGATATCTTAAGAGGAAgaTGTTATTCGCTAACACCTGCAC |
| KBP1 | myo2-F1264E | CTCATTTGTGGTGgaaGCTCTAAACTCTATTTTAACCG |
| KBP2 | myo2-F1275E | CCGAAGAAACGgaaAAAAACGGCATGA |
| LHP352 | myo2-L1411W | GTCGTGGAAAAGGGGTtggCAATTGAACTACAACG |
| LHP353 | myo2-T1418Q | GGGGTCTTCAATTGAACTACAACGTTcaaAGATTAGAGGAATGG |
| LHP356 | myo2-I1458A | CGAAGACATTGATgcCTTAAGAGGAATTTGTTATTCGC |
| LHP357 | myo2-E1484R | GGTGGCAGACTATagGTCTCCAATTCCAC |
| LHP344 | myo2-Y1483A | AGGTGGCAGACgcTGAGTCTCCAA |
| LHP347 | myo2-K1408A | GTAATTTCTTGTCGTGGgcAAGGGGTCTTC |
| LHP348 | myo2-E1338R | TTTCAGCGCGGGAcgAACCAGCGGGTTTT |

| | | |
|---|---|---|
| LHP325 | vac17-F132Y | ccagaagtagtttagggtcatAtcaacctcgacc |
| LHP326 | vac17-L137A | catttcaacctcgaccaGCgaaaataattgagaggc |
| LHP328 | vac17-R142E | ccattgaaaataattgagGAgcaacgtctgtgtatggtaactcc |
| LHP253 | vac17-K138E | gtcatttcaacctcgaccattgGAaataattgagaggcaacg |
| LHP255 | vac17-R135E | gggtcatttcaacctGAaccattgaaaataattgagaggc |
| **Internal deletions of Vac17** | | |
| LHP339 | Q5SDM_12/2/2022_R 110- | GGCCAGCTCGTTCAACCT |
| LHP340 | Q5SDM_12/2/2022_F -157 | GTAGGCTTTAACCCCATCAATG |
| LHP107 | Q5SDM_HHPRED pRS416-vac17(18-108 del)-3XENVY_F | GCCACCGTCCCAAATGAA |
| LHP108 | Q5SDM_HHPRED pRS416-vac17(18-108 del)-3XENVY_R | CGACCTTATTAAAAGCCTCTC |

**Table S3 Strains used in this study**

| Strain | Genotype | Reference |
|---|---|---|
| LSW5798 | *MATa*, *ura3-52*, *leu2-3,-112*, *his3-Δ200*, *trp1-Δ901*, *lys2-801*, *suc2-Δ9*, *vac17Δ::TRP1* | Tang *et al*., 2003 |
| LWS8195 | *MATα, ura3-52, leu2-3,-112, his3-Δ200, trp1-Δ901, lys2-801, suc2-Δ9, pep4-Δ1137, vac17Δ::TRP1, myo2Δ::TRP1, YCp50-MYO2* | Yau *et al*., 2014 |
| LSW9506 | *MATa, vac17Δ, vac8Δ, myo2Δ::YCP50-MYO2* | Yui Jin |
| LSW13976 | *MATa, vac17Δ, VPH1-mCherry* | Yui Jin |
| LSW8226 | *MATa, YFP-PTS1::LEU2, myo2 delta::TRP1, YCP50-MYO2* | Yui Jin |
| LSW7656 | *MATa, leu2,3-112 ura3-52 his3-Δ200 trp1-Δ901 lys2-801 suc2-Δ9 vac8Δ::VAC8-mRFP-HIS3 vac17Δ::TRP1* | Tang *et al*., 2006 |

**Table S4 Plasmids used in this study**

| Plasmid name | Description | Reference |
|---|---|---|
| pRS416-Vac17 | CEN, URA3 | Catlett *et al*., 1998 |
| pRS416-Vac17-3xEnvy | CEN, URA3 | This study |
| pRS416-*vac17Δ18-108*-3xEnvy | CEN, URA3 | This study |
| pRS416-*vac17Δ195-250*-3xEnvy | CEN, URA3 | This study |
| pRS416-*vac17Δ110-157*-3xEnvy | CEN, URA3 | This study |
| pRS413-Myo2 | CEN, HIS3 | Yau *et al*., 2013 |
| pRS413-mCherry-Myo2 | CEN, HIS3 | Jin *et al*., 2011 |
| pRSF-Duet-1 | Kan | Gift from Amir Kahn |
| pRSF-Strep II-Myo2 Tail(1150-1567) | Kan | This study |
| pMALc2H10T | Amp | Kristelly *et al*., 2008 |
| pMBP-10xhis-TEV-Vac17(1-157) | Amp | This study |
| pMBP-10xhis-TEV-Vac17(110-157) | Amp | This study |
| pMBP-10xhis-TEV-Vac17(1-109) | Amp | This study |

**Table S5 Cryo-EM Data Collection, Refinement, and Validation Statistics**

| | Vac17+Myo2 tail complex<br>EMD |
|---|---|
| **Data collection** | |
| Grids | UltrAuFoil |
| Vitrification method | FEI Vitrobot |
| Automation software | Leginon |
| Session name | 23jun08b |
| Microscope | Titan Krios G3 |
| Voltage (keV) | 300 |
| Recording mode | Counting |
| Detector | K3 |
| Energy filter slit width | Gatan Bioquantum imaging filter (20 eV) |
| Magnification | 81,000X |
| Pixel size at detector (Å/pixel) | 1.085 |
| Total electron exposure (e–/Å$^2$) | 73.2 |
| Exposure rate (e-/pixel/sec) | 21.5 |
| Number of frames | 40 |
| Defocus range (µm) | 0.8-2.0 |
| **Data processing** | |
| Software(s) used | CryoSPARC v4 |
| Micrographs collected (no.) | 4,605 |
| Micrographs used (no.) | 3,224 |
| Particles extracted (no.) | 1,387,051 |
| Final particle images (no.) | 618,699 |
| Final particles in refinement | 144,724 & 31,238 |
| Symmetry | C1 |
| Map resolution (Å) | 5.75 & 7.21 |
| FSC 0.5 | 7.4/6.3/5.8 & 8.1/7.0/7.3 |
| (unmasked/masked/corrected) | -585.7 & -451.8 |
| Map sharpening B factor | |
| **Refinement** | |
| Refinement package | Real-space refinement on PHENIX (v1.20.1-4487) |
| Initial model used | Trimmed Alphafold model of Vac17(127-147) bound Myo2(1151-1569) |
| **Model composition** | |
| Non-hydrogen atoms | 3,430 |
| Protein residues | 423 |
| **Validation** | |
| MolProbity score | 1.74 |
| CaBLAM outliers | 0.24 |
| **Ramachandran plot** | |
| Favored (%) | 98.32 |
| Allowed (%) | 1.68 |

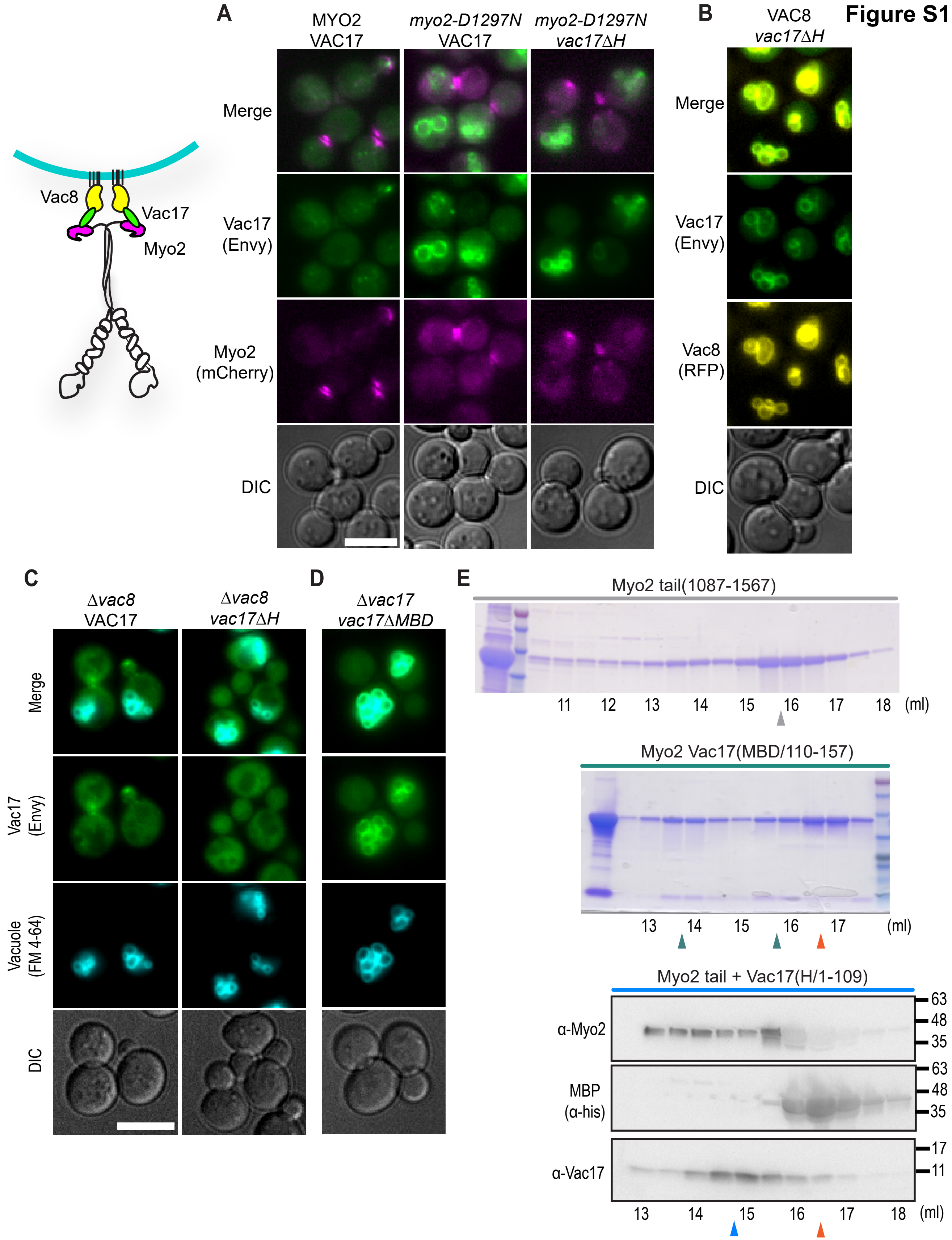
Figure S1
A
MYO2
VAC17
myo2-D1297N
VAC17
myo2-D1297N
vac17ΔH
Merge
Vac17
(Envy)
Myo2
(mCherry)
DIC
Vac8
Vac17
Myo2
B
VAC8
vac17ΔH
Merge
Vac17
(Envy)
Vac8
(RFP)
DIC
C
Δvac8
VAC17
Δvac8
vac17ΔH
D
Δvac17
vac17ΔMBD
Merge
Vac17
(Envy)
Vacuole
(FM 4-64)
DIC
E
Myo2 tail(1087-1567)
11
12
13
14
15
16
17
18
(ml)
Myo2 Vac17(MBD/110-157)
13
14
15
16
17
(ml)
Myo2 tail + Vac17(H/1-109)
α-Myo2
MBP
(α-his)
α-Vac17
63
48
35
63
48
35
17
11
13
14
15
16
17
18
(ml)

Figure S2

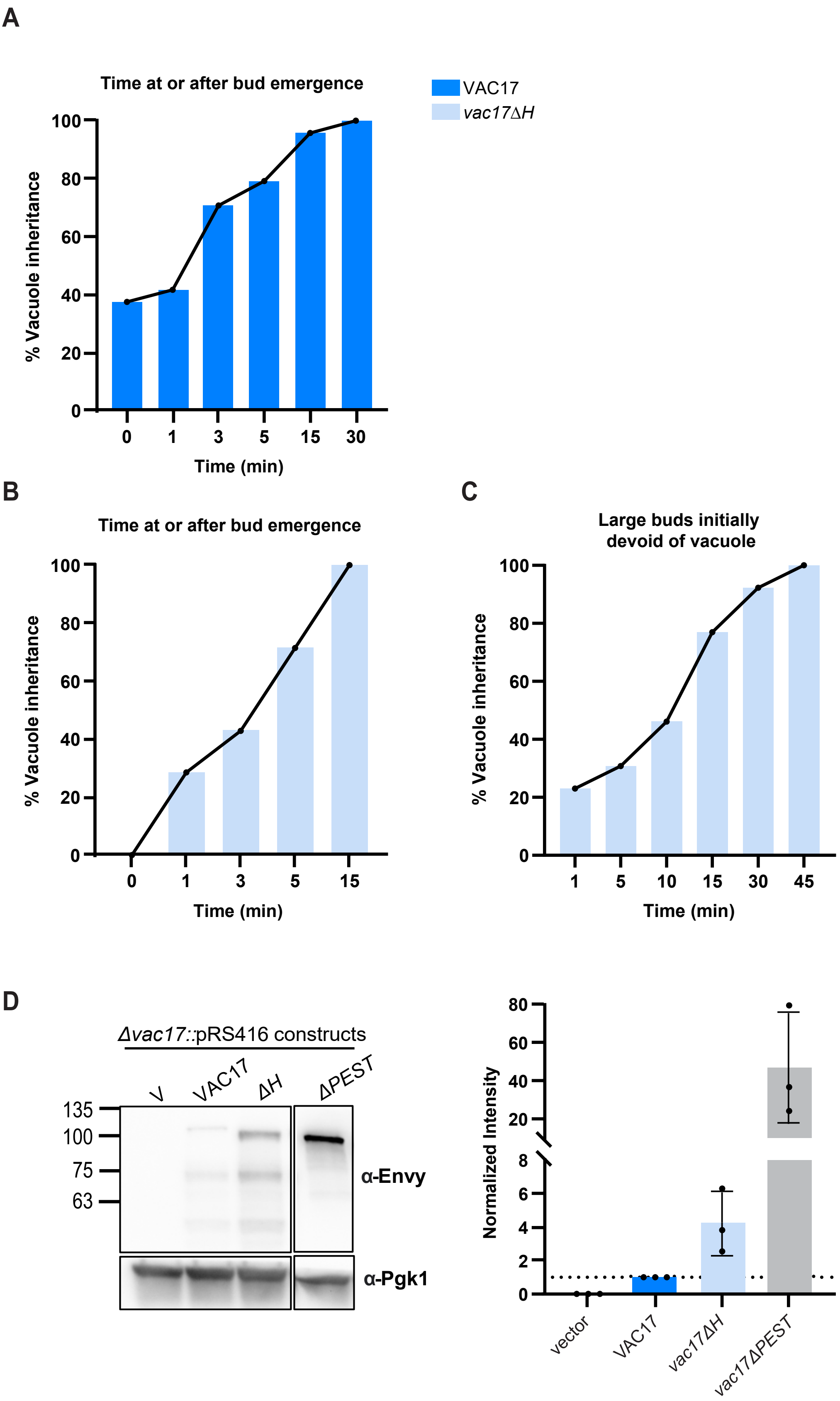
A
Time at or after bud emergence
VAC17
vac17ΔH
% Vacuole inheritance
Time (min)
B
Time at or after bud emergence
% Vacuole inheritance
Time (min)
C
Large buds initially devoid of vacuole
% Vacuole inheritance
Time (min)
D
Δvac17::pRS416 constructs
V
VAC17
ΔH
ΔPEST
135
100
75
63
α-Envy
α-Pgk1
Normalized Intensity
vector
VAC17
vac17ΔH
vac17ΔPEST

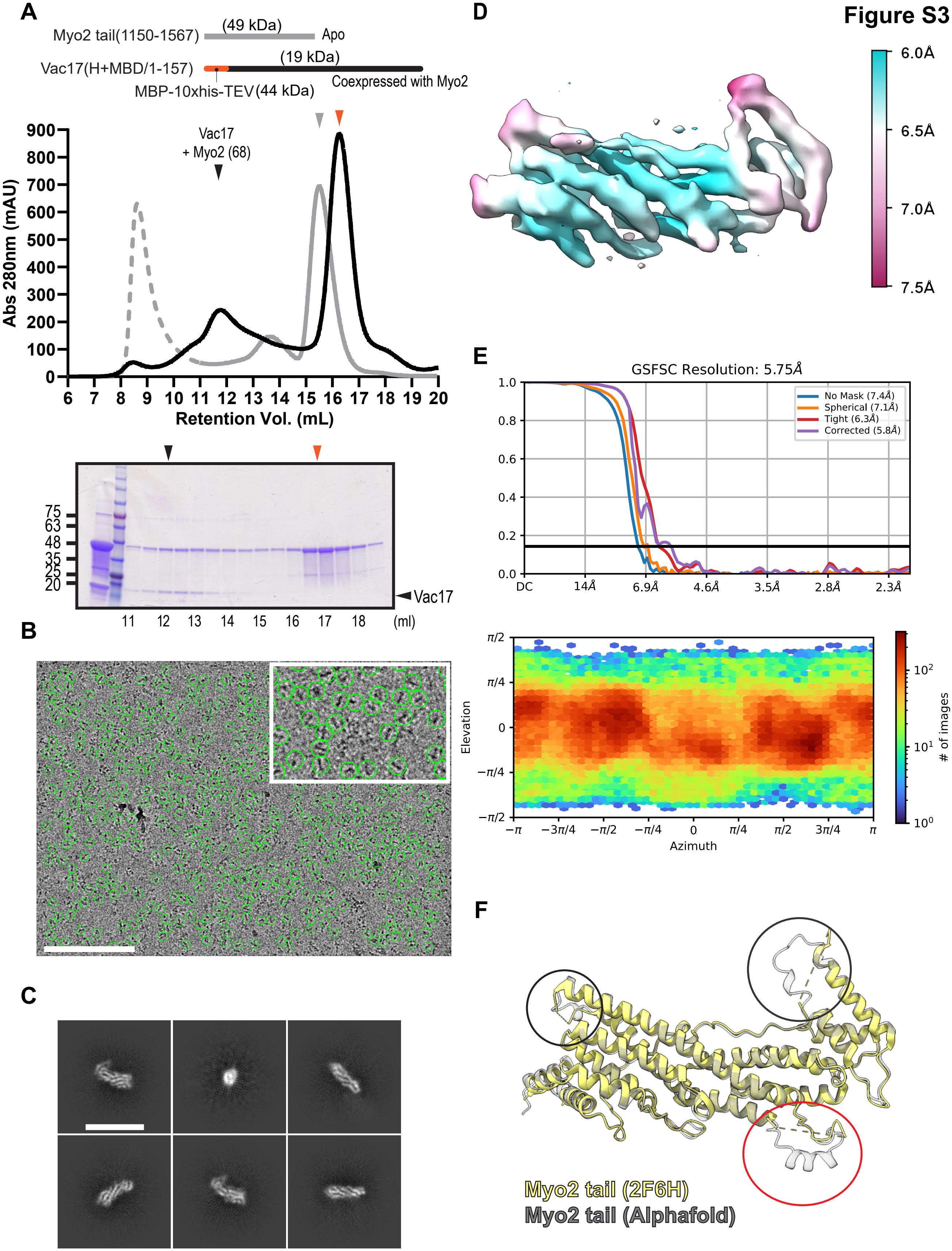

A
Myo2 tail(1150-1567)
(49 kDa)
Apo
Vac17(H+MBD/1-157)
(19 kDa)
MBP-10xhis-TEV
(44 kDa)
Coexpressed with Myo2
Vac17 + Myo2 (68)
Abs 280nm (mAU)
Retention Vol. (mL)
75
63
48
35
25
20
Vac17
11 12 13 14 15 16 17 18 (ml)
B
C
D
6.0Å
6.5Å
7.0Å
7.5Å
E
GSFSC Resolution: 5.75Å
No Mask (7.4Å)
Spherical (7.1Å)
Tight (6.3Å)
Corrected (5.8Å)
DC
14Å
6.9Å
4.6Å
3.5Å
2.8Å
2.3Å
Elevation
Azimuth
### of images
F
Myo2 tail (2F6H)
Myo2 tail (Alphafold)

Figure S3

Figure S4

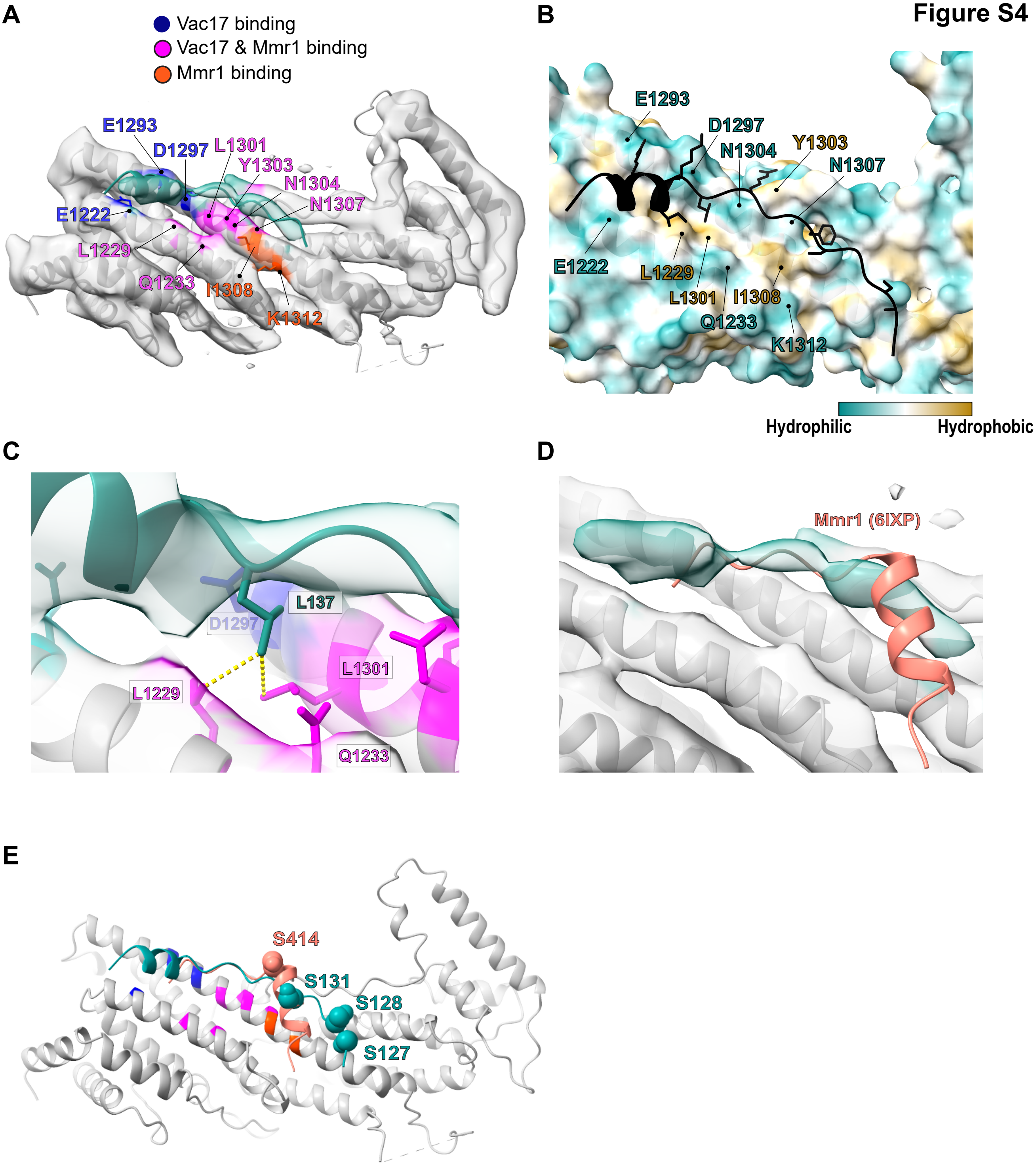
A
Vac17 binding
Vac17 & Mmr1 binding
Mmr1 binding
E1293
D1297
L1301
Y1303
N1304
N1307
E1222
L1229
Q1233
I1308
K1312
B
E1293
D1297
N1304
Y1303
N1307
E1222
L1229
L1301
I1308
Q1233
K1312
Hydrophilic
Hydrophobic
C
L137
D1297
L1301
L1229
Q1233
D
Mmr1 (6IXP)
E
S414
S131
S128
S127

**Figure S5**

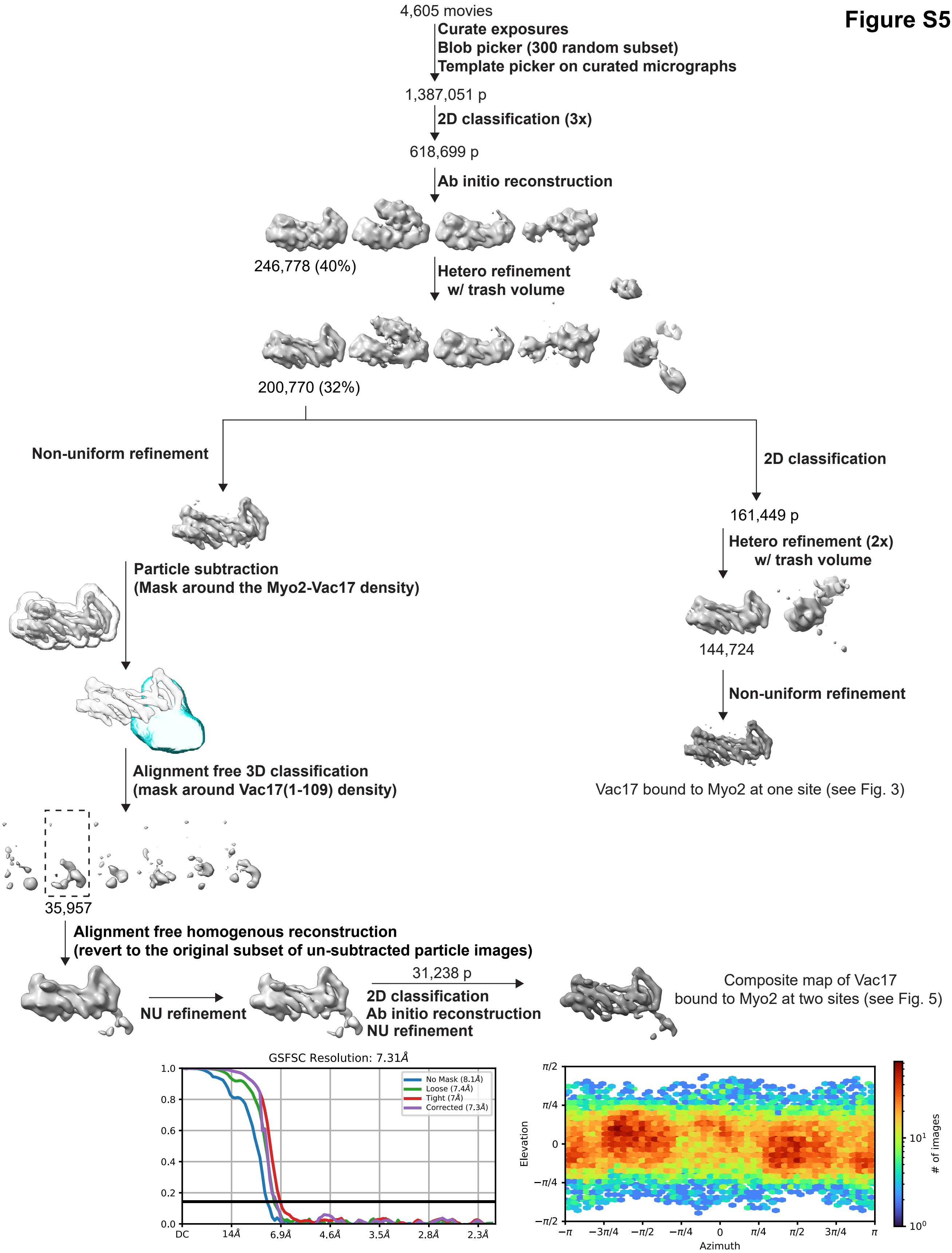
4,605 movies
Curate exposures
Blob picker (300 random subset)
Template picker on curated micrographs
1,387,051 p
2D classification (3x)
618,699 p
Ab initio reconstruction
246,778 (40%)
Hetero refinement
w/ trash volume
200,770 (32%)
Non-uniform refinement
2D classification
161,449 p
Hetero refinement (2x)
w/ trash volume
144,724
Non-uniform refinement
Vac17 bound to Myo2 at one site (see Fig. 3)
Particle subtraction
(Mask around the Myo2-Vac17 density)
Alignment free 3D classification
(mask around Vac17(1-109) density)
35,957
Alignment free homogenous reconstruction
(revert to the original subset of un-subtracted particle images)
NU refinement
31,238 p
2D classification
Ab initio reconstruction
NU refinement
Composite map of Vac17
bound to Myo2 at two sites (see Fig. 5)
GSFSC Resolution: 7.31Å
No Mask (8.1Å)
Loose (7.4Å)
Tight (7Å)
Corrected (7.3Å)
1.0
0.8
0.6
0.4
0.2
0.0
DC
14Å
6.9Å
4.6Å
3.5Å
2.8Å
2.3Å
π/2
π/4
0
−π/4
−π/2
Elevation
−π
−3π/4
−π/2
−π/4
0
π/4
π/2
3π/4
π
Azimuth
10^1
10^0
### of images
